# Evolutionary Dynamics of RuBisCO: Emergence of the Small Subunit and its Impact Through Time

**DOI:** 10.1101/2024.06.06.597628

**Authors:** Kaustubh Amritkar, Bruno Cuevas-Zuviría, Betül Kaçar

**Affiliations:** Department of Bacteriology, University of Wisconsin-Madison, Madison, WI, USA; Biophysics Graduate Degree Program, University of Wisconsin-Madison, Madison, WI, USA; Centro de Biotecnología y Genómica de Plantas, Universidad Politécnica de Madrid, Madrid, Spain

**Keywords:** RuBisCO, small subunit, ancestral sequence reconstruction, structural dynamics

## Abstract

Ribulose-1,5-bisphosphate carboxylase/oxygenase (RuBisCO) is an ancient protein critical for CO_2_-fixation and global biogeochemistry. Form-I RuBisCO complexes uniquely harbor small subunits that form a hexadecameric complex together with their large subunits. The small subunit protein is thought to have significantly contributed to RuBisCO’s response to the atmospheric rise of O_2_ ∼2.5 billion years ago, marking a pivotal point in the enzyme’s evolutionary history. Here, we performed a comprehensive evolutionary analysis of extant and ancestral RuBisCO sequences and structures to explore the impact of the small subunit’s earliest integration on the molecular dynamics of the overall complex. Our simulations suggest that the small subunit restricted the conformational flexibility of the large subunit early in its history, impacting the evolutionary trajectory of the Form-I RuBisCO complex. Molecular dynamics investigations of CO_2_ and O_2_ gas distribution around predicted ancient RuBisCO complexes suggest that a proposed “CO_2_ reservoir” role for the small subunit is not conserved throughout the enzyme’s evolutionary history. The evolutionary and biophysical response of RuBisCO to changing atmospheric conditions on ancient Earth showcase multi-level and trackable responses of enzymes to environmental shifts over long timescales.

## Introduction

Life has been evolving on this planet for four billion years. Evolution allows for exploration of the vast sequence space comprised by polymers of the twenty standard amino acids – a space that exceeds the number of molecules in the universe (Wagner and Rosen 2014). From within this space, biology has discovered unique solutions to the challenges of growing and persisting in the ever-shifting diversity of Earth’s environments. Undoubtedly, biogeochemically critical microbial enzymes are central to this dynamic, long-term interaction between life and the environment (Falkowski et al. 2008). One such key enzyme that evolved early in the history of life and persisted through planetary extremes is RuBisCO (ribulose-1,5-bisphosphate carboxylase/oxygenase).

RuBisCO is a globally critical, ancient enzyme with an intriguing history. It facilitates the rate-limiting step of the Calvin-Benson-Bassham cycle for carbon fixation, catalyzing the addition of atmospheric carbon dioxide (CO_2_) with ribulose 1,5-bisphosphate (RuBP) (Andersson 2008). In the presence of oxygen (O_2_), RuBisCO also catalyzes a competing oxygenation reaction in which RuBP combines with O_2_, producing an autoinhibitory metabolite detrimental to the overall metabolic efficiency of carbon fixation (Fernie and Bauwe 2020). While the exact age of RuBisCO is not clearly known, paleobiological inferences (Ward and Shih 2019; Garcia et al. 2021; Kędzior et al. 2022) suggest that it emerged prior to the rise of atmospheric O_2_ ∼2.5 billion years ago, a period known as the Great Oxidation Event (GOE). Accordingly, despite being infamously sensitive to O_2_, RuBisCO was maintained by organisms through this drastic atmospheric upheaval that impacted the ecosystem and its constituent biomolecules (Ashida et al. 2005; Raymond and Segrè 2006; Wang et al. 2011; Young et al. 2012; Caetano-Anollés 2017; Kacar et al. 2017; Erb and Zarzycki 2018; Garcia et al. 2021).

Extant RuBisCO forms (I to IV) exhibit different dimeric and poly-dimeric assemblies. All forms have large subunits (RbcL), but Form-I RuBisCOs uniquely have an additional small subunit (RbcS) (**Fig 1A**). Form-I RuBisCOs are also the most abundant (Tabita et al. 2008) and generally exhibit higher specificities for CO_2_ (Flamholz et al. 2019). These attributes make RbcS a promising candidate to study how small accessory subunits can regulate the evolution of RuBisCO, though their precise evolutionary role remains unclear. RbcS is thought to be crucial for the assembly of the RbcL octameric complex made of RbcL homodimers (Liu et al. 2010; Esquivel et al. 2013; Joshi et al. 2015) and significantly impacts RuBisCO’s catalytic parameters (Gatenby 1988; Spreitzer 2003; Genkov et al. 2010; Esquivel et al. 2013; Matsumura et al. 2020; Mao et al. 2022). For example, incorporation of ancestral RbcS in RuBisCO was shown to increase the enzyme’s specificity for CO_2_, indicating that emergence of RbcS played a critical role in the ancestor of Form-I RuBisCO (Schulz et al. 2022). Previous computational work also suggested that RbcS can act as a “CO_2_-reservoir” to concentrate CO_2_ molecules within the enzyme (Van Lun et al. 2014). However, the extent to which the small subunit has influenced the structural dynamics and evolutionary trajectory of the Form-I RuBisCO complex remains less understood.

**Fig. 1:**
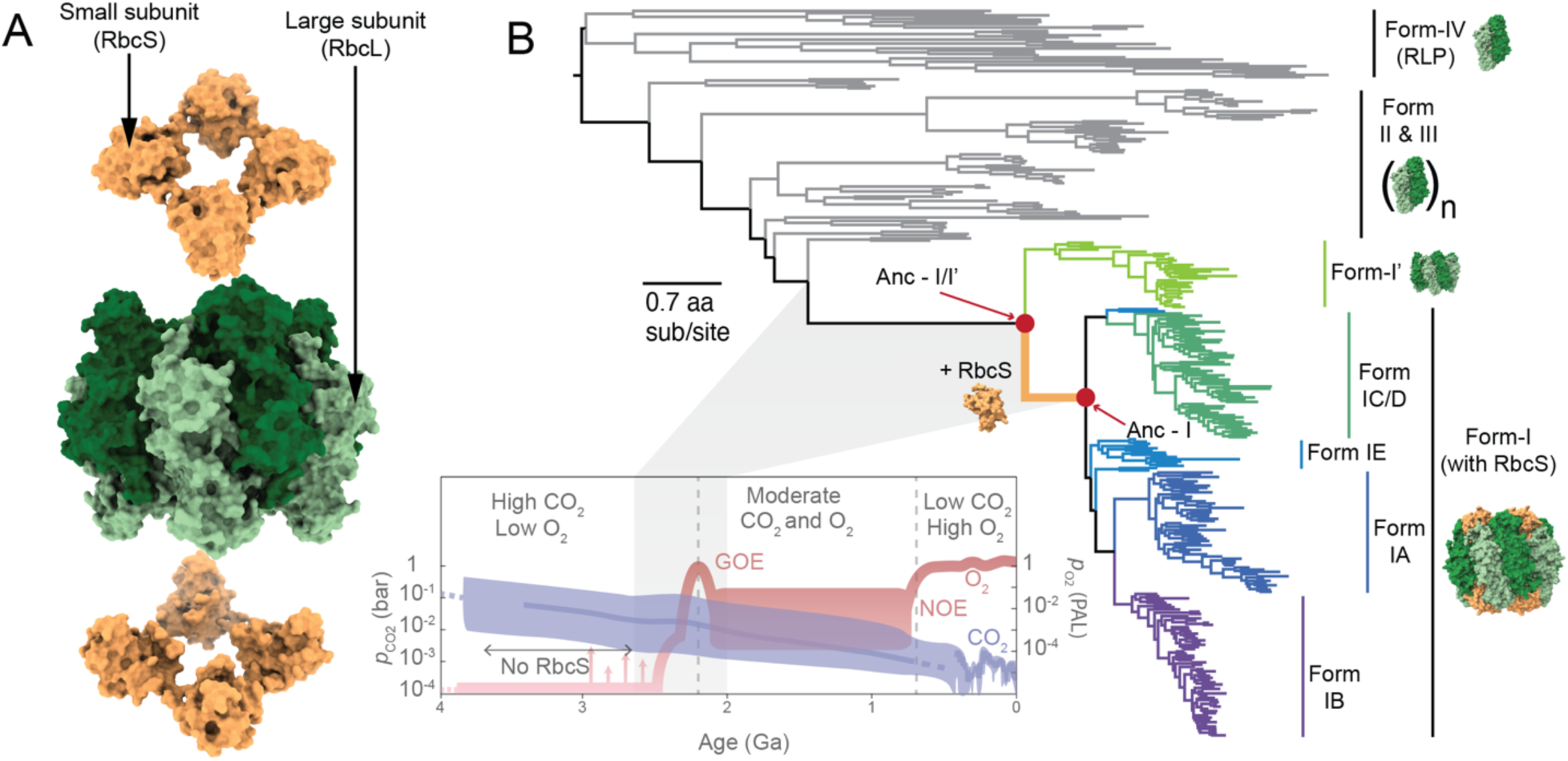
The evolutionary history and diversification of RuBisCO. (A) Architecture of the L_8_S_8_ structure of the Form-I RuBisCO complex, with RbcL shown in green and RbcS in orange (PDB:1BWV). (B) RuBisCO phylogenetic tree highlighting the emergence of RbcS coinciding with the Great Oxidation Event (GOE) (Kacar et al. 2017; Banda et al. 2020). Schematic on the right shows variation in the multimeric structure of the RuBisCO complex across different forms. Form-II and III RuBisCOs exhibit multiple homooligomeric states of the RbcL- dimer, such as RbcL-dimer, tetramer or hexamer (Liu et al. 2022), represented here as (RbcL- RbcL)_n_, where n is the number of RbcL-dimers. Bottom inset schematic shows the change in atmospheric CO_2_ and O_2_ concentration through Earth’s history (Rucker and Kaçar 2023).

Here we used phylogenetic reconstructions, structural predictions as well as molecular dynamics (MD) simulations to track the evolution of the small subunit in the context of the RuBisCO complex over geologic time. Our exploration focused on the hypothesis that the molecular and structural innovations likely made Form-I RuBisCO selectively more advantageous following the planetary rise of O_2_. We specifically focused on three areas: 1) the sequence and structural evolution of subunits within ancient RuBisCO complexes following integration of ancient RbcS, 2) the impact of RbcS presence or absence on the structural dynamics of ancestral and extant RuBisCO complexes, and 3) migration of CO_2_ and O_2_ gases in ancestral RuBisCO to assess a proposed CO_2_-reservoir role for ancient small subunits (Van Lun et al. 2014).

## Results and Discussion

### Resurrection of ancestral RuBisCOs

We built a maximum-likelihood phylogenetic tree from a concatenated alignment of RbcL and RbcS amino acid sequences representing RuBisCO Forms I to IV, including the recently described Form-I’ (Banda et al. 2020) (**Fig. 1B**). The phylogeny contains 194 sets of RbcL-RbcS homologs from Form-I and 135 RbcL homologs from all other forms (including Form-I’, II, II/III, III and IV). The tree contains sequences representative of known RuBisCO diversity and is rooted by Form-IV RuBisCO-like proteins in accordance with previous studies (Tabita et al. 2007; Kacar et al. 2017; Poudel et al. 2020). Form-I is categorized into four major subgroups: the “green-like” Form-IA and IB, prevalent in proteobacteria, cyanobacteria, green algae and plants, and the “red-like” Form-IC and ID, prevalent in proteobacteria and non-green algae (Spreitzer 2003; Tabita et al. 2008). Our phylogeny also resolves a Form-I subclade, Form-IE, that diverges before the last common ancestor of Form-IA and Form-IB (West-Roberts et al. 2021). Most sequences from the Form-IE clade are from metagenomic studies and belong to unclassified members of the *Chloroflexota* bacterial phylum. Form-I’ RuBisCOs cluster within a monophyletic clade sister to all other Form-I sequences. Form-I’ RuBisCOs are notable because their RbcL subunits are similar to those of Form-I in their multimeric assembly, but they lack the RbcS (Banda et al. 2020). Like Form-IE, the Form-I’ clade contains sequences from *Chloroflexota*. The topology of our concatenated phylogeny is in agreement with previously reported RbcL-based phylogenetic trees (Kacar et al. 2017; Banda et al. 2020).

We selected ancestral nodes situated along the evolutionary trajectory immediately before and after RbcS incorporation into RuBisCO for ancestral sequence reconstruction (Methods). Specifically, we inferred the common ancestor of both Form-I’ and Form-I RuBisCOs, as well as the ancestors of major Form-I clades as highlighted in **Fig. 2A**, inferring a total of seven ancestral nodes along the phylogenetic tree.

**Fig. 2:**
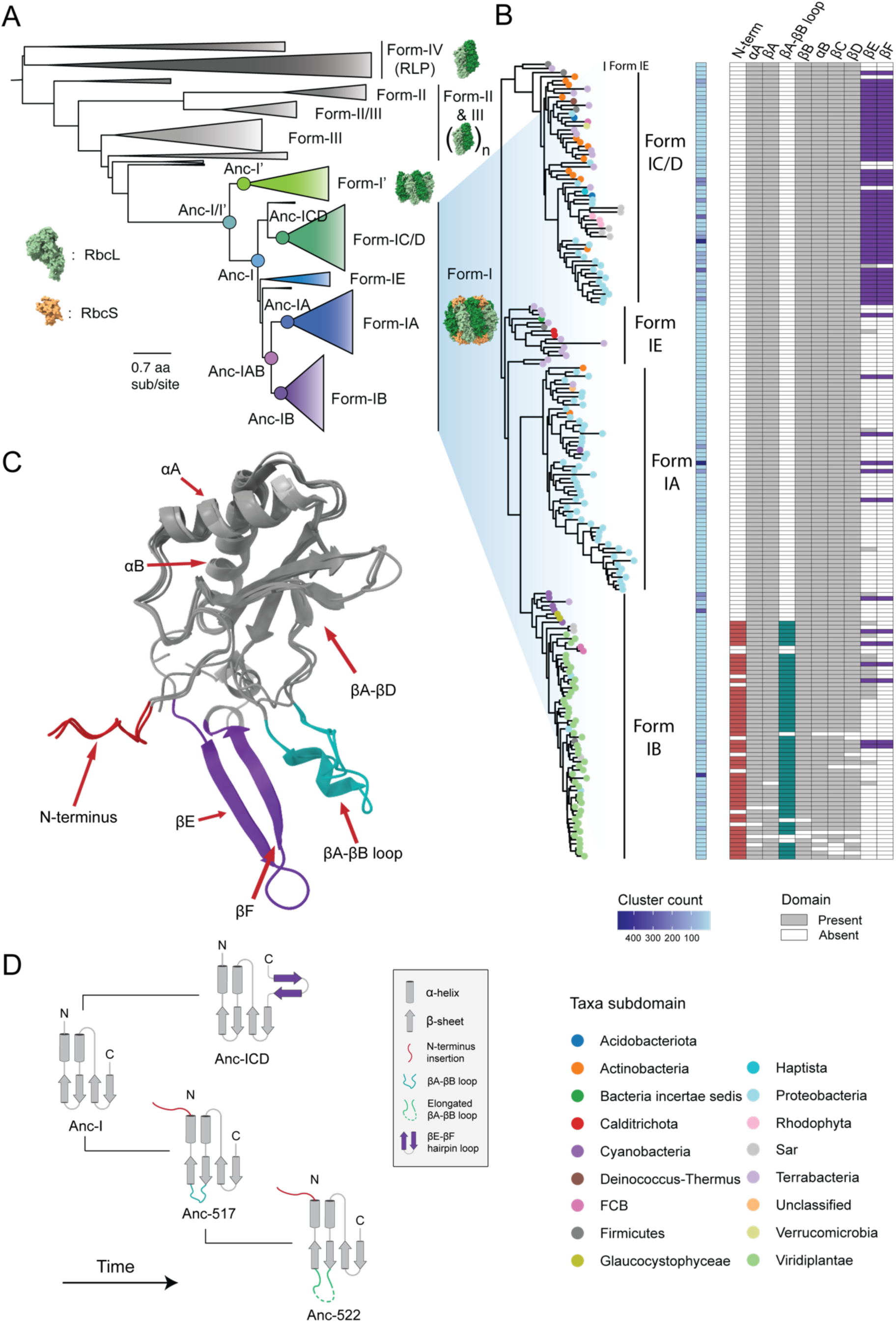
Phylogenetic analysis of RbcS structural diversity. (A) Collapsed maximum-likelihood phylogenetic tree of concatenated RbcL-RbcS sequences. Ancestral nodes used in this study are highlighted. (B) Analysis of RbcS structural diversity, mapped to a clustered Form-I phylogeny, where each tip represents a cluster of RbcS sequences with >63% sequence identity. A heatmap for the number of sequences present in the RbcS cluster is represented in blue. Extant nodes are colored by microbial host taxonomic diversity. The presence and absence of different RbcS features in each extant sequence cluster is indicated by gray and white colors, and the color scheme for the presence of uncommon features is according to (C). (C) Multiple RbcS structures across Form-I subgroups aligned to highlight the different RbcS structural features. (D) Schematic representing the emergence of RbcS structural features across the Form-I RuBisCO phylogeny.

### Sequence and structural diversity of modern and ancient RbcS proteins

We analyzed all extant Form-I RbcS sequences from our database representative of known host taxonomic diversity (**Fig. 2B**, Methods). Extant Form-I RbcS homologs show remarkably low mean pairwise sequence identity (29%) compared to Form-I RbcL sequences (66%) (**supplementary fig. 3**) as observed by others previously (Bracher et al. 2017; Mao et al. 2022; Bouvier et al. 2024).

We inferred sequences for 328 ancestral nodes within the phylogenetic tree, with 193 of these nodes belonging to the Form-I clade, including its common ancestor, Anc-I. Form-I RbcL ancestors are reconstructed with higher confidence than Form-I RbcS ancestors, with mean posterior probabilities of 0.98 and 0.90, respectively, across all positions (**supplementary fig. 4**). Anc-I subunit sequences have a mean pairwise identity of 71.5% (RbcL) and 42.2% (RbcS) compared across all extant RuBisCOs in the phylogeny, indicating a higher conservation of RbcL compared to RbcS. The mean posterior probabilities for Anc-I RbcL and RbcS are 0.95 and 0.85 respectively.

We further explored historical, evolutionary variation of RbcS by identifying known, functionally significant sequence and structural features (Spreitzer 2003; Mao et al. 2022) (**Fig. 2C**) and mapping their presence across extant and reconstructed ancestral RbcS in our phylogenetic tree (**Fig. 2B and 2D**). We find that the RbcS structure contains novel features unique to different Form-I specific lineages. While all RuBisCO small subunits share a common core structure consisting of a ꞵ-sheet with four antiparallel strands (ꞵA to ꞵD) and two α-helices (αA & αB) (Knight et al. 1990), the Form-IB and Form-IC/D RbcS exhibit additional distinct characteristics.

Within the Form-IB clade, we find that most green-like eukaryotic RbcS homologs contain an additional N-terminal region as well as a ꞵA-ꞵB loop insertion. The N-terminal region is specifically known to be a signal peptide necessary for the entry of RbcS into the chloroplast, prior to assembly with RbcL (Schmidt and Mishkind 1986; Spreitzer 2003). The N-terminal extension feature is first observed in the Form-IB ancestry after the separation of cyanobacteria in the phylogeny (**supplementary fig. 5**). Absence of the N-terminal extension in Anc-IB, the common ancestor of Form-IB, as well as older ancestors indicates that their ancient hosts did not translate RbcS and RbcL in separate organelles. Rather, expression of subunits in different organelles emerged as a trait following the bifurcation of cyanobacteria and Viridiplantae in Form-IB RuBisCO.

We find that the ꞵA-ꞵB loop insertion is similarly exclusive to green-like eukaryotic RbcS (**Fig. 2B**). Although the significance of the ꞵA-ꞵB loop in RbcS is not established, it plays an important part in regulating the size of the central solvent channel in RuBisCO complex (Esquivel et al. 2013). Like N-terminal extension, this loop first emerges after the splitting of cyanobacterial and eukaryotic lineages in Form-IB (**supplementary fig. 5**). As observed with extant RbcS (Spreitzer 2003; Mao et al. 2022), the ꞵA-ꞵB loop insertion length also varies across Form-IB ancestors (10 to 28 aa). The loop has increased by ∼11 residues in all eukaryotic ancestors after the divergence from cyanobacterial ancestors and by ∼18 residues in the ancestors from green algae lineage within Form-IB (**supplementary fig. 5**).

We also tracked the evolution of the C-terminal ꞵE and ꞵF hairpin loop that is unique to Form-ICD RbcS sequences (**Fig. 2B**). The common ancestor of Form-IC/D sequences, Anc-ICD, and all subsequent Form-IC/D ancestors have the ꞵE and ꞵF hairpin (**Fig. 2D**). By contrast, Anc-I RbcS does not have the ꞵE and ꞵF hairpin, suggesting this insertion happened after the divergence of Form-IC/D within the Form-I clade. The C-terminal ꞵ-hairpin is functionally significant for mediating the assembly of the oligomeric complex, allowing red-like RuBisCOs to assemble without specialized assembly chaperones (Joshi et al. 2015). The incorporation of the hairpin loop in Form-IC/D common ancestor, Anc-IC/D, suggests that independence from assembly chaperones may have first emerged in a red-like RuBisCO ancestor. Our analysis thus reveals that RbcS has undergone different clade-specific structural variations since its emergence in the Form-I ancestor.

### Global distribution of residues evolutionarily linked to RbcS incorporation

RbcL from Form-I and Form-I’ RuBisCOs exhibit the same L_8_ oligomeric arrangement. Because the presence or absence of the small subunit is the only major structural distinction between the two forms (Banda et al. 2020), we performed comparative analyses to reveal the small subunit’s impact on RbcL sequence and structure. We focused on the RbcL extant sequences from the Form-I and Form-I’ clades. We identified residues that are conserved within each clade, but differ between the two clades, which we refer to as “signature positions”. These residues are more likely to contribute to the functional distinctions between Form-I and Form-I’, including interactions (or lack thereof) with RbcS. The interface region between the large and small subunit has been shown to be critical for the incorporation of RbcS into the Form-I RuBisCO complex (Knight et al. 1990; Van Lun et al. 2011; Ryan et al. 2019; Schulz et al. 2022). We therefore hypothesized that signature positions would cluster within the RbcL-RbcS interface.

Form-I and Form-I’ sequence analyses revealed eleven, discontinuous signature positions (**Fig. 3A**). These positions are clustered in three different regions on the RuBisCO structure. Two residues (Val 70 and Asp 74; position index from *Rhodobacter sphaeroides*) are located at the RbcL intradimer interface (highlighted in blue), three (Phe 201, Leu 401 and Gln 402) are buried in the RbcL monomer close to the active site (highlighted in purple) and six (Lys 260, Trp 284, Arg 286, Asn 288, Gly 159 and Glu162) are found at the RbcL dimer-dimer interface (highlighted in red) (**Fig. 3A**). Out of the six residues located at the RbcL dimer-dimer interface, Gly 159 and Glu 162 are the only two signature positions that are also present at the RbcL-RbcS interface. A similar comparison between the Form-IA/B/E and Form-IC/D RbcS sequences did not identify any signature positions (Methods).

**Fig. 3:**
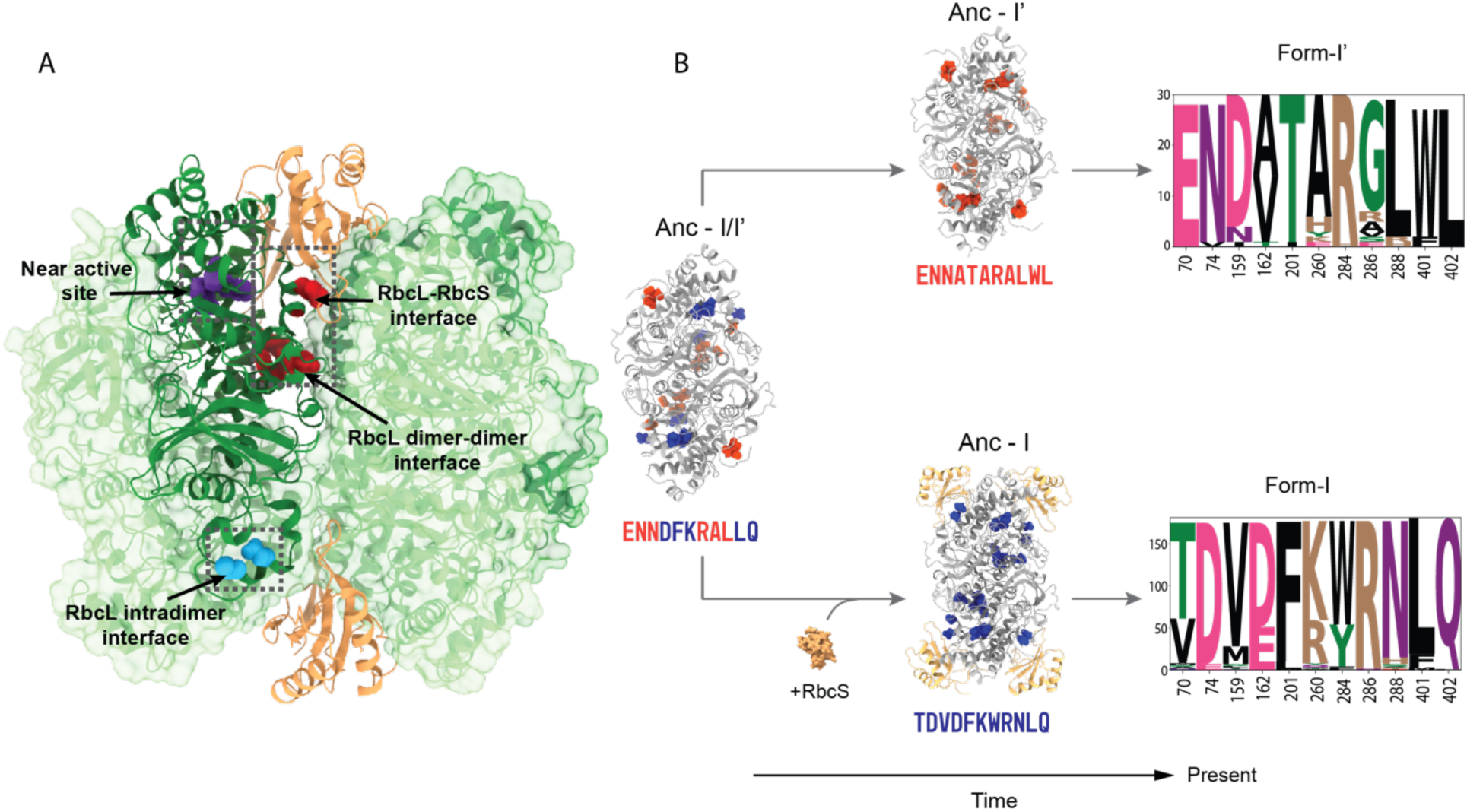
Schematic representation of the structural distribution and evolution of the specialized protein “signature positions” between the RuBisCO Form-I and Form-I’. (A) Signature residues are highlighted based on their structural proximity to each other along with the region it belongs to on the *Rhodobacter sphaeroides* L_4_S_2_ structure (PDB: 5NV3). Residues at the signature positions located at the RbcL intradimer interface, near active site and RbcL dimer-dimer interface are highlighted in blue, purple and red respectively. (B) Evolution of the signature residues corresponding to the divergence of Form-I and Form-I’ RuBisCOs. Amino acids present in the sequence motif are highlighted on the respective ancestor’s L_2_ or L_2_S_4_ structures. Sequence logos showing amino acid frequencies at each signature position across the Form-I and Form-I’ clades. Residue numbering according to the *R. sphaeroides*, L denotes RbcL and S denotes RbcS.

We examined how these signature positions evolved through the emergence of RbcS by tracking their ancestral amino acid composition through the divergence of Form-I and Form-I’ clades. Residues at the signature positions in Anc-I and Anc-I’ most closely resemble those of their respective descendants (**Fig. 3B**). By contrast, the common ancestor of all Form-I and Form-I’ RuBisCOs, Anc-I/I’, has approximately equal proportions of Form-I- and Form-I’-like signature positions. Among these, signature positions at the Anc-I/I’ RbcL intradimer interface (labeled 70,74) are Form-I’-like, whereas those near the RbcL active site (labeled 201, 401, 402) are Form-I-like. Notably, the Anc-I/I’ RbcL dimer-dimer interface and RbcL-RbcS interface contains a mixture of Form-I- and Form-I’-like signature positions. These results identify the interface residues that are likely important for oligomerization of the RuBisCO complex and underwent distinct functional specialization in the presence (in Form-I) or absence (in Form-I’) of RbcS.

In sum, the signature positions are not restricted to the RbcL-RbcS interface but are scattered across the RbcL structure (**Fig. 3**). We suggest that RbcS integration resulted in distant mutations, which are known to influence protein function by altering intramolecular interactions and the global dynamics of protein complexes (Miton et al. 2021). We therefore submit that these positions reflect how the presence or absence of RbcS affected long-range motions and conformational dynamics of the RuBisCO complex, and therefore its evolutionary dynamics.

### Molecular dynamics simulations of modern and ancient RuBisCOs

We studied the structural motions of RuBisCO complexes to investigate the influence of the presence or absence of RbcS. We built the ancestral RuBisCO large and small subunit structures (Methods). Modeling of the multimeric structural assembly of ancestral RuBisCO oligomers was guided by extant RuBisCO crystal structures (Methods). We simulated the molecular dynamics of seven ancestral and six extant RuBisCO complexes representing a range of host organisms and their associated environments (**Fig. 4A**, **Table 1**). We performed two simulation replicates for each RuBisCO variant, and data from both replicates are included in the analysis.

**Fig. 4:**
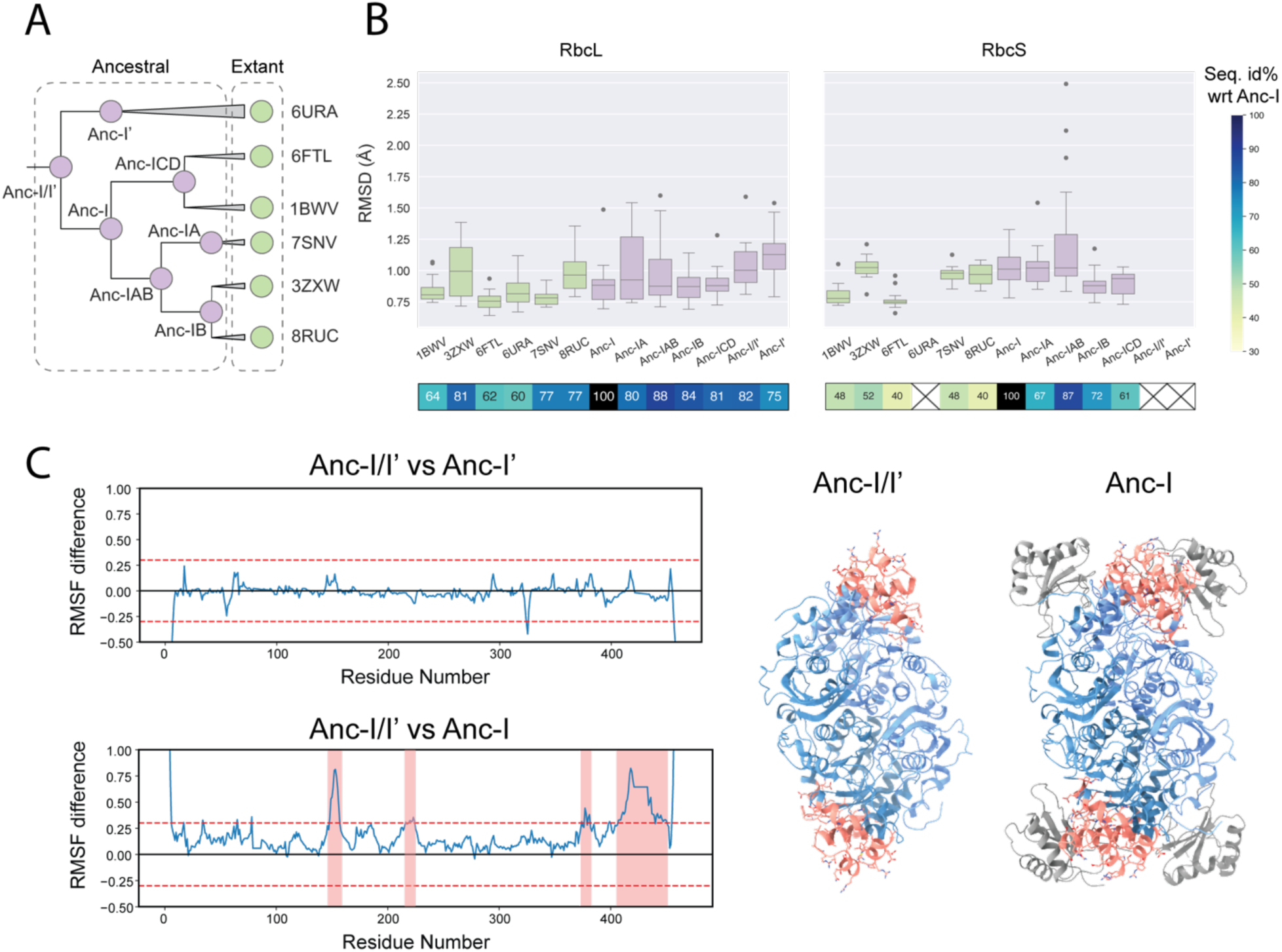
Stability and fluctuation observed in the ancestral and extant RuBisCOs during MD simulations. (A) Ancestral and extant RuBisCO homologs utilized in the study. Violet circles: ancestral RuBisCO, green circle: extant RuBisCO. (B) Average root mean square deviation (RMSD) of all amino acids in the large and small subunits throughout the simulations. Outliers outside the quartile range are represented by circles. Heatmap on the x-axis represents the pairwise sequence identity for each sequence relative to the oldest RbcS ancestor, Anc-I. (C) Difference in residue-wise root mean square fluctuation (RMSF) for RbcL between Anc-I/I’ vs Anc-I’ (top), and Anc-I/I’ vs Anc-I (bottom). Horizontal dashed red line represents the RMSF difference of 0.3 Å, used as threshold to classify the residue fluctuation as considerable, and the horizontal black line represents the baseline when there is a zero difference in RMSF between the residues from the two proteins. Residues with RMSF over 0.3 Å are highlighted in red on the Anc-I/I’ and Anc-I L_2_ and L_2_S_4_ complex structures respectively on the right.

**Table 1:**
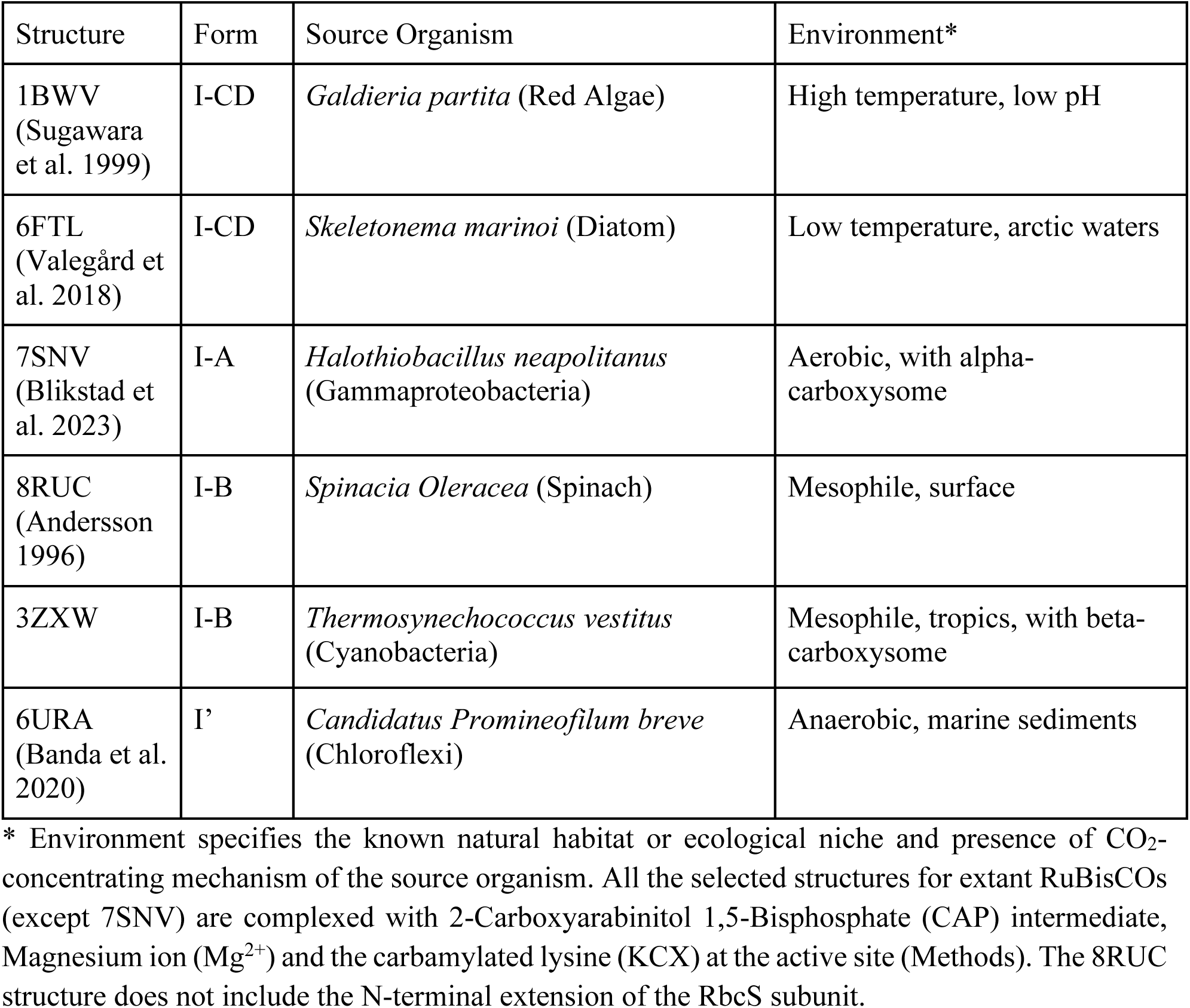
Overview of extant RuBisCO structures selected for analysis.

We calculated the root mean square deviation (RMSD) of backbone atoms for all residues in the RbcL and RbcS subunits relative to their average trajectory structure to evaluate conformational differences in the complex during the simulation. The mean RMSD of all the individual large and small subunits across all complexes is 0.928Å and 0.964Å, respectively. The average RbcL RMSD ranges from 0.78Å to 1.13Å, while the RbcS RMSD ranges from 0.76 to 1.26Å (**Fig. 4B)**. The outliers in Anc-IAB small subunit RMSD values (**Fig. 4B)** reflect the N-terminal residues residue protrusion and unfolding observed across the simulation replicates. We performed pairwise comparisons of mean RMSD values among the different RuBisCOs. Ten out of all possible pairs show a significant pairwise RMSD difference for RbcL (p < 0.01, Tukey post-hoc test). And eight of these include the Anc-I’ or Anc-I/I’ complex, with a significantly higher RMSD in each case (**supplementary table 3**). A higher RMSD for the Anc-I/I’ and Anc-I’ RbcL suggests a greater structural flexibility compared to the other complexes during the simulations.

We used the root mean square fluctuation (RMSF) of the different residues to identify region-specific differences in flexibility. Specifically, we studied the ancestral structures immediately before and after divergence of RbcS-less Form-I’ and RbcS-containing Form-I RuBisCOs. These ancestors include Anc-I/I’, Anc-I’, and Anc-I. The alteration in residue-wise fluctuations between the Anc-I’ and Anc-I/I’ complex during the MD simulation is close to zero (**Fig. 4C, top**). In contrast, the Anc-I RbcL exhibits less fluctuation compared to Anc-I/I’ during the simulations, where four distinct sections from the large subunit sequence show considerably higher fluctuations Anc-I/I’ compared to Anc-I (**Fig. 4C, bottom**). The sections with higher flexibility correspond to the top region of the large subunit that is positioned between the two small subunits in the RuBisCO complex (**Fig. 4C, right)**. Out of the 45 RbcL residues present at the RbcS interface in Anc-I, 23 residues (∼51%) show a significant difference in RMSF. The localization of RbcL residues near RbcS suggests that the presence of RbcS restricts the movement of these residues. This observation potentially accounts for the higher stability of the Anc-I large subunit compared to the RbcS-less Anc-I/I’ and Anc-I’. The differences in RMSF lend support to the hypothesis that RbcS plays a role in stabilizing the RbcL dimers in the octameric complex (Spreitzer 2003).

The extant Form-I’ RuBisCO (6URA) provides an additional basis for comparative analysis. Along with Anc-I’ and Anc-I/I’, RbcS is also absent in 6URA. Unlike its ancestor, however, the extant Form-I’ complex does not display a significantly higher RMSD during the pairwise comparison to the other RuBisCO systems (**supplementary table 3**). Similarly, 6URA RbcL displays lower fluctuations per residue compared to the Anc-I/I’ and Anc-I’ RbcL (**supplementary fig. 9**). This indicates that in the absence of RbcS, the modern Form-I’ RbcL has evolved a more stable L_8_ without RbcS, potentially through other mutations. The L_8_ assembly in the extant Form-I’ complex structure is maintained by a network of hydrogen bonds and salt bridges (Banda et al. 2020). Some of the key residues Asp161, Trp165 and Tyr224 involved in these interactions in the extant 6URA structure are not conserved in the analyzed RbcS-less ancestors, Anc-I’ and Anc-I/I’ (Asn140, Arg144 and Phe203 respectively) (**supplementary fig. 7**). The absence of these key residue interactions may explain the higher fluctuations observed in the Anc-I’ and Anc-I/I’ RbcLs during the simulation.

While RbcS is indispensable for the activity of extant Form-I RuBisCOs (Andrews 1988; Lee and Tabita 1990), its present-day importance does not necessarily imply that it played an essential role at its initial emergence. Protein subunits can become entrenched in complexes via accumulations of neutral mutations that can be deleterious in monomers (Hochberg et al. 2020). However, our analysis further indicates the early importance of RbcS for enhanced stability of the RbcL octamer, along with prior experimental work demonstrating enhanced specificity towards CO_2_ (Schulz et al. 2022). These findings collectively suggest an early advantage of RbcS in RuBisCO assembly at the time of its acquirement.

### Impact of RbcS integration on the conformational variation of Form-I RbcL

The global distribution of RbcL signature positions (**Fig. 3**), inferred to be functionally linked to the presence of RbcS, suggests that RbcS integration likely affected the large subunit’s overall dynamics and motion. We utilized MD simulations for RuBisCO complexes with and without RbcS to investigate the impact of RbcS on the enzyme’s ability to explore conformational space. A protein’s conformational space can be visualized as a set of different conformational states and the conformational variability of an enzyme is considered as an important factor for its functional diversity (Nobeli et al. 2009; Babtie et al. 2010).

To characterize the variation in conformations for the different RuBisCO simulations, we performed a principal component analysis (PCA) of the trajectory of the RbcL Cα atoms (Hayward and de Groot 2008; David and Jacobs 2014). This analysis allows us to reduce the dimensionality of the simulation trajectories into a few relevant dimensions, referred to as principal components (PC). To compare different RbcLs, we extracted the coordinates of conserved residues across all RbcLs and projected the trajectories onto a shared set of PCs (Methods). The dimensionality reduction shows the changes along the top two PCs (PC1 and PC2) for the RbcL’s MD trajectories, capturing ∼27% of all RbcL motions during the simulations (**Fig. 5A**). We observe that RbcL from each RuBisCO variant clusters closely together (**Fig. 5B**) suggesting that the dynamics of each individual RbcL are similar to those of its counterparts within the RuBisCO complex.

**Fig. 5:**
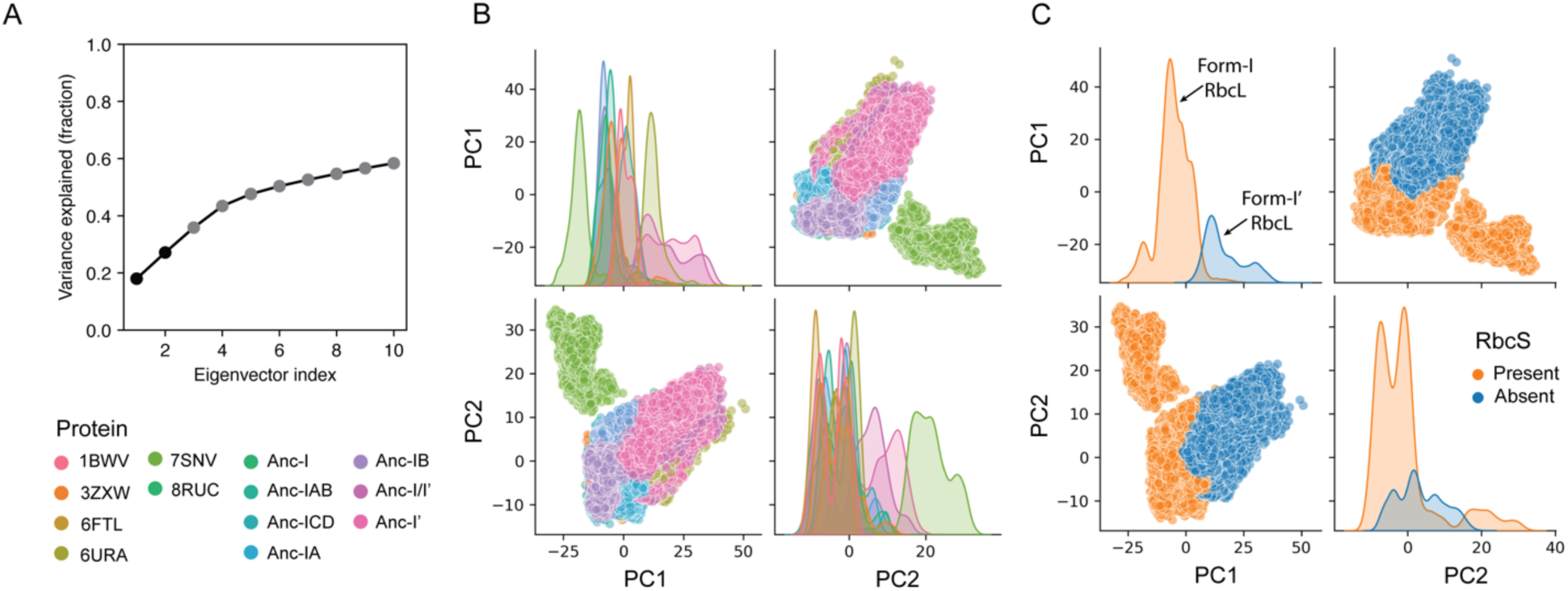
Principal Component Analysis for the conserved RbcL residues of MD-simulation trajectories across ancestral and extant Form-I and Form-I’ RuBisCO variants. (A) Variance explained by each eigenvector in the PCA, showing the contribution of the top 10 principal components to overall fluctuation across all simulations. (B) Pairwise representation of PC1 and PC2, highlighting the different Form-I and Form-I’ RuBisCO variants. (C) Pairwise representation of PC1 and PC2, highlighting the RbcS presence and absence in the RuBisCO complex.

We observe that Form-I RuBisCOs (with RbcS) and non-Form-I RuBisCOs (without RbcS) occupy distinct regions along PC1 (**Fig. 5C**) with statistical significance (U-statistic = 2.09×10^7^; p = 0.0; Mann-Whitney U test). In contrast, PC2, does not show a specific trend between RuBisCO variants. This suggests that the primary conformational variation (∼19% of all RbcL motions) corresponds to the distinction between Form-I and non-Form-I RuBisCOs.

We simulated the dynamics of Anc-I, the common ancestor of Form-I RuBisCO, without the RbcS subunit and projected its trajectory onto the previously defined PCs for Form-I and Form-I’. The results show an intermediate positioning between Form-I and Form-I’ RuBisCOs (**supplementary figure 10A**). Removing RbcS from Form-I RuBisCO, shifts it dynamics closer to Form-I’, but does not fully replicate Form I’ behavior. This indicates that RbcS plays a critical, though not exclusive, role in driving the distinct conformational variations between Form-I and Form-I’.

We assessed the impact of octameric oligomerization of RbcL on enzyme dynamics by including MD simulations of a Form-II RuBisCO (9RUB) in our PCA decomposition. Incorporating Form-II RbcL generates a distinct set of PCs (*PC’*). The first PC (PC1’) separates Form-II from Form-I and Form-I’; while the second PC (PC2’) distinguishes Form-I and Form-I’ (**supplementary figure 10B**). This separation suggests that the octamerization in Form-I/Form-I’ induces significant conformational changes compared to the non-octameric Form-II, with the Form-I and Form-I’ distinction becoming apparent only in PC2’.

Our results suggest that the presence or absence of RbcS in the RuBisCO complex impacts the major dynamics of RbcL. We hypothesize that the incorporation of RbcS induces a conformational shift in RbcL, a modification that has remained consistent across diverse Form-I and Form-I’ RuBisCO variants over evolutionary time. The functional diversity (e.g., promiscuity) of a protein is closely tied to its ability to explore diverse conformational states (Tokuriki and Tawfik 2009; Zou et al. 2015; Petrović et al. 2018; Jackson et al. 2022). In this context, our analysis shows that the RbcL explores distinct conformations in the presence of RbcS, suggesting that the shift in RuBisCO’s conformational variability may be directly associated with the enzyme’s increased CO_2_ specificity following the emergence of RbcS (Schulz et al. 2022).

### Heterogeneity of CO_2_/O_2_ diffusion patterns across ancestral and extant RuBisCOs

Previous work suggested that RbcS acts as a CO_2_-reservoir (Van Lun et al. 2014), increasing the concentration of CO_2_ molecules near the enzyme and making CO_2_ more accessible to the active site. This hypothesis also provides an additional functional justification for the integration of RbcS, given the increase in Earth’s atmospheric O_2_ concentration ∼2.5 billion years ago, around the time of RbcS integration into the RuBisCO complex.

We performed MD simulations for the different RuBisCO complexes with CO_2_ and O_2_ gas molecules present in the medium (**Fig. 6A**) to assess the interaction of RbcS residues towards both gases. We estimated the relative contact-scores of the gas molecules by counting the number of interactions between the gases and the protein throughout the simulation within a distance threshold of 6Å. The MD simulations for O_2_ and CO_2_ gas molecules were performed separately (**Fig. 6A**). Five independent replicates for MD simulations of one extant (8RUC) and one ancestral (Anc-I) enzyme were conducted to assess the robustness and replicability of the gas dynamics modeling approach. To assess the impact of gas concentrations, we performed simulations with varying CO_2_ and O_2_ gas concentrations. The trends observed for affinity of the subunits across different gas concentrations remains relatively consistent between the two subunits (**supplementary fig. 11)**.

**Fig. 6:**
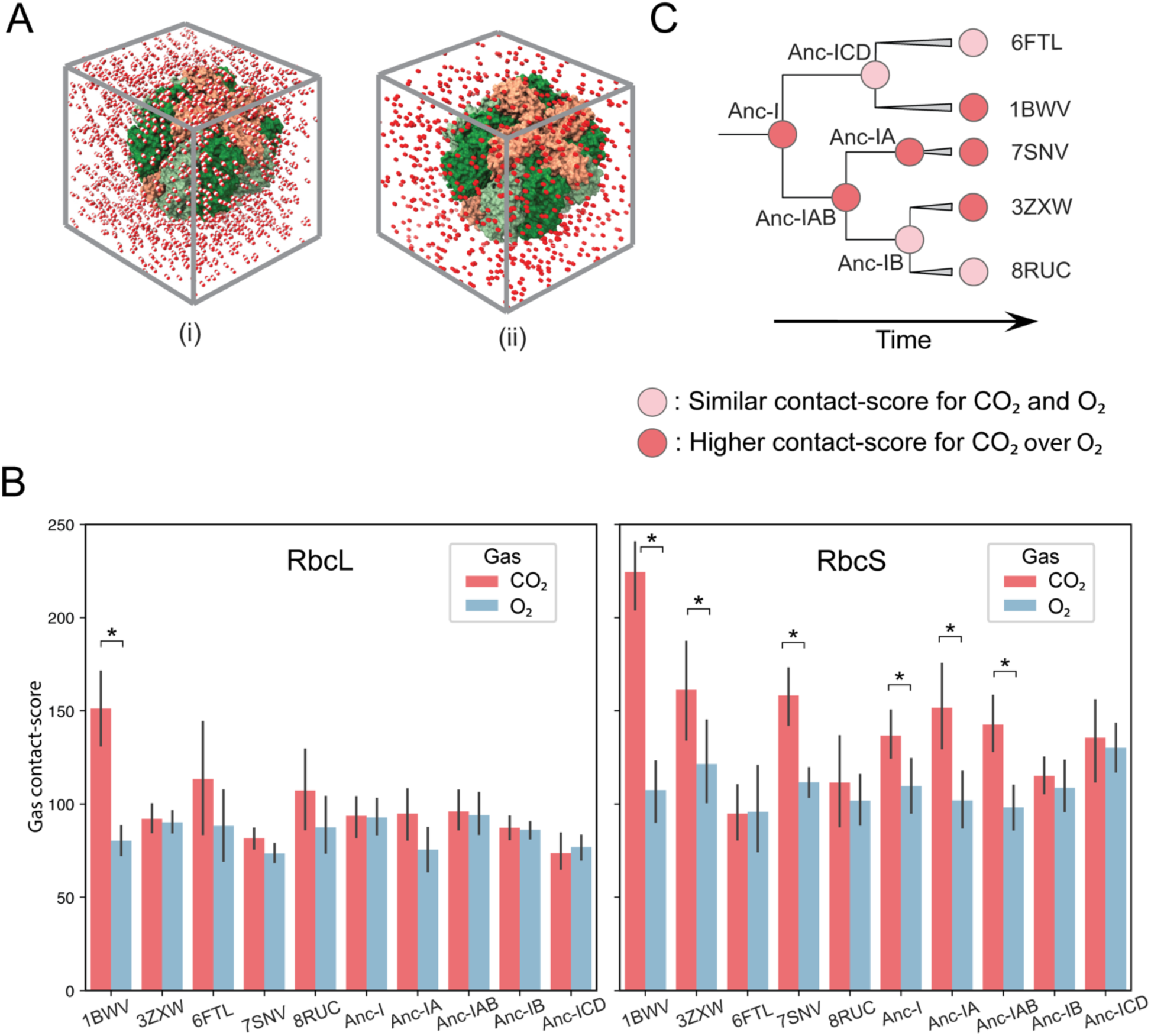
CO_2_ and O_2_ gas diffusion molecular dynamics across different RuBisCO systems. (A) A snapshot of the MD simulation box with CO_2_ (i) and O_2_ (ii) gas molecules in the medium at a concentration of 400 mM with the protein complex. (B) Barplot representing the contact-score (number of gas contacts per residue) for RbcL and RbcS for each of RuBisCO for the two CO_2_ and O_2_ gas molecules. Error bars represent the standard deviation across the 8 subunits in the RuBisCO complex. Proteins with a significant difference between the CO_2_ and O_2_ affinity are marked with an asterisk (*). (C) Trend for the difference in contact-score for CO_2_ and O_2_ gas molecules for RbcS as observed in the barplot (B). Time is displayed along the y-axis.

An independent two sample t-test was performed to compare the mean CO_2_ and O_2_ gas contacts across all simulations. For RbcL, we observe that only 1BWV has a significantly higher contact-score for CO_2_ over O_2_ (t = 5.92; p = 3.7×10^-5^, two-sided), whereas other complexes do not show a statistically significant (p > 0.05) difference (**Fig. 6B**). Alternatively, six RbcS (1BWV, 3ZXW, 7SNV, Anc-I, Anc-IA, Anc-IAB) out of the ten exhibit a significantly (p < 0.05) higher contact-score for CO_2_ than for O_2_ molecules in the surrounding medium (**Fig. 6B**, **supplementary table 4**). RbcS that exhibit a higher affinity for CO_2_ over O_2_ are not confined to a specific clade in the phylogenetic tree (**Fig. 6C**). We find that these results do not vary significantly over five independent simulations (**supplementary fig. 12)**.

## Conclusions

The co-evolution of the large and small subunits of RuBisCO presents an opportunity to analyze molecular evolutionary events within key macroevolutionary and geochemical transitions, independently documented by Earth’s geologic record. We examined the historical impact of RuBisCO small subunit emergence on the structural motions and evolutionary trajectory of the enzyme, with specific focus on a possible adaptive role in concentrating CO_2_. Both extant and ancestral Form-I RuBisCO complexes that contain RbcS show higher CO_2_-specificity than other forms of RuBisCO lacking RbcS (Flamholz et al. 2019; Schulz et al. 2022). These observations have led to the hypothesis that the improved CO_2_-specificity of RuBisCO after RbcS integration would have been advantageous after the GOE, given the subsequent rise in atmospheric O_2_ levels and decrease in CO_2_ concentrations. We illustrate that not all ancestral and extant RbcS proteins act as a reservoir to concentrate CO_2_ around the RuBisCO complex. It is possible that certain RuBisCOs with RbcS might have adopted a CO_2_-reservoir strategy based on other intercellular or possibly environmental factors as we find that this feature appears multiple times in the enzyme’s history. Moreover, our analysis highlights additional consequences of RbcS integration: increased stability of Form-I RbcL and decreased flexibility compared to Form I’ RbcL. The small subunit integration may have shaped the enzyme’s major conformational and functional variations in response to significant shifts in atmospheric composition.

The origin of the small subunit still remains unresolved, but these observations shed light on the circumstances that surrounded and likely facilitated the emergence of RbcS. Certain cyanobacteria have two other proteins that are part of the carbon fixation machinery: ꞵ- carboxysome structural protein (CcmM) and a RuBisCO activase-like protein (ALC). Both these proteins have one or multiple domains that are “RbcS-like” and are considered homologs to RbcS (Ryan et al. 2019; Wang et al. 2019; Lechno-Yossef et al. 2020). The presence of these protein domains is specific to cyanobacteria. Further studies should explore whether the RbcS-like protein domains could have emerged alongside or following the RbcS in response to the ancient shifts in atmospheric CO_2_ and O_2_ levels. In summary, we show that ancestral RuBisCO dynamically responded to a global environmental shift. The integration of the small subunit resulted in increased rigidity, allowing it to maintain substrate specificity despite decreased substrate availability. Ironically, this natural solution to an ancient challenge may have led to modern RuBisCO’s notorious resistance to artificial improvements in CO₂ specificity.

## Methods

### Phylogenetic Reconstruction of RuBisCO

Homologous sequences for RbcS protein from the NCBI non-redundant database were identified using PSI-BLAST (Altschul et al. 1997; Altschul and Koonin 1998). The RbcS sequence from *Thermosynechococcus elongatus* (PDB id: 2YBV) (Gubernator et al. 2008) was used as the query sequence for 5 PSI-BLAST iterations with 50,000 sequences per iteration and an E-value cutoff of 0.005. The dataset was curated to remove partial sequences and sequences with incomplete annotations. The sequences in the dataset were dereplicated at 63% amino acid identity with CD-Hit (Li and Godzik 2006) and aligned by MAFFT (Katoh et al. 2002) (default parameters) with iteration refinement over 1000 cycles. Any poorly aligned sequences were subsequently replaced with a different representative of their CD-HIT cluster.

A maximum-likelihood phylogenetic tree was constructed using IQ-Tree (Nguyen et al. 2015) with the LG+R6 evolutionary model (**supplementary fig. 1**). ModelFinder Plus (Kalyaanamoorthy et al. 2017) was used to select the best-fit evolutionary model. Branch support values were calculated using the Shimodaira–Hasegawa–like approximate likelihood-ratio test (SH-aLRT) with 1000 bootstrap replicates and 1000 ultrafast bootstrap (UFBoot) replicates optimized by nearest neighbor interchange (NNI).

RbcL sequences from the same taxa represented in our RbcS sequence dataset were identified using the NCBI Identical Protein Groups database (IPG) (https://www.ncbi.nlm.nih.gov/ipg). For RbcS entries with missing IPG RbcL sequences, Taxonomy-restricted BLASTp was used to search for the corresponding RbcL sequence. Form-I RbcL sequences were combined with RbcL sequences for Form-I’, II, III and IV from Banda et al. (Banda et al. 2020) to build an RbcL dataset. An RbcL phylogenetic tree was constructed using IQ-Tree with the LG+R9 evolutionary model (all other parameters were the same as for the RbcS tree) (**supplementary fig. 2**).

The sequences in the RbcL dataset were aligned by MAFFT, with the same parameters as those mentioned above, and were concatenated with the RbcS alignment. A maximum-likelihood phylogenetic tree was constructed for the concatenated alignment using IQ-Tree, with a partition model (Chernomor et al. 2016) and ModelFinder plus (Kalyaanamoorthy et al. 2017) to search and implement the best-fit evolutionary models corresponding to the RbcL (LG + R9) and RbcS (LG + R5) section of the concatenated alignment. In line with prior research, Form-IV “RuBisCO-like” protein sequences were used to root the phylogenetic tree (Kacar et al. 2017; Banda et al. 2020; Poudel et al. 2020; Camel and Zolla 2021; Schulz et al. 2022). Ancestral sequence inference for the concatenated RbcL-RbcS tree was performed using PamL4.9 (LG model) (Yang 2007). Reconstruction of the gaps in the ancestral sequences was performed using a binary likelihood model as described by Aadland et al. (Aadland et al. 2019).

### RbcS sequence and structural diversity analysis

Candidate RbcS structures (listed in **Table 1**) from Form-I subclades were used to identify the different structural features in RbcS. The amino acid regions corresponding to these features were identified in the multiple sequence alignment of extant RbcS. The binary heatmap indicating the presence or absence of these features across all extant RbcS sequences in the phylogeny was generated based on the presence of at least half of the residues in the respective region of the RbcS alignment. Host taxonomy for the extant sequences was assigned using the REST API provided by Ensembl (https://rest.ensembl.org/).

### RbcL specialized signature position analysis

Sequence specialization between the Form-I and Form-I’ extant RbcL sequences was analyzed using TwinCons (Penev et al. 2021), with blosum62 as substitution matrix for score calculation, and Zebra2 (Suplatov et al. 2020), using the webserver default parameters. Specialized residues identified by Zebra2 and TwinCons score were then filtered based on conservation in the multiple sequence alignment for Form-I and Form-I’ sequences. Same steps as above were followed to identify separately conserved amino acids between 131 RbcS sequences from Form-IA/B/E and 59 from Form-IC/D. None of the RbcS positions had a TwinCons score < −1, resulting in no signature positions.

### Structural Modeling of RuBisCO

We predicted the structures of ancestral RuBisCOs using the deep-learning based Colabfold software (Mirdita et al. 2022). We built our models by combining predictions and structure alignments. We built the model for the RbcL-dimer with RbcS-dimer for each ancestral protein with templates from PDB, structure refinement using Amber and 3 prediction recycles. Low-confidence modeled terminal regions for the predicted structures were removed using a pLDDT threshold of 50. The extant RuBisCO structure from *Thermosynechococcus elongatus* (PDB: 2YBV) was used as a template to obtain a predicted hexadecameric complex using UCSF Chimera (Pettersen et al. 2004). Anc-I’ and Anc-I/I’ did not have an RbcS sequence resulting in an octameric complex.

### RuBisCO MD simulations

We simulated the molecular dynamics of both extant and ancient RuBisCO complexes. We selected seven ancestral structures (nodes highlighted in **Fig. 2A**) corresponding to the last common ancestors of different Form-I and Form-I’ clades and six experimentally determined extant RuBisCO structures, representing the Form-IA, IB (two from this clade), IC/D (two from this clade), and I’ RuBisCO clades. PDB IDs for these entries are: 1BWV (red algae), 3ZXW (green algae), 6FTL (diatom), 6URA (chloroflexi), 7SNV (gammaproteobacteria) and 8RUC (plantae) (more details in **Table 1**). The 6URA entry represents the first case discovered of a Form-I-like RuBisCO without a small subunit. This selection represents the biological diversity of the RuBisCO enzyme while preserving the following attributes: high resolution (under 3.0 Å), carbamylation of the lysine at the active site (except for 7SNV) (Stec 2012), and CABP and magnesium at the active site. Sequence identity and alignments for ancestral and extant RuBisCOs analyzed in the study are presented in **supplementary fig. 6 and 7**, respectively.

We obtained parameters for the CABP molecule through the standard Amber protocol for ligands (Wang et al. 2004). RuBisCO proteins include different post-translational modifications that required additional steps. We computed the CABP and the carbamylated lysine charges at HF/6-31*G level, as specified in the Amber protocol. The post-translational modifications found in 6FTL were modeled with semi-empirical charges. In the case of the modeled ancient structures, we manually added both ligands (CABP), ions (Mg^2+^), and the post-translation modification at the carbamylated lysine (also for 7SNV).

We built the topology and parameters of the molecular dynamics systems using tleap (Maier et al. 2015) and Amber14 (Salomon-Ferrer et al. 2013). Periodic solvation boxes were constructed with 10 Å spacing and water molecules according to the TIP3P model (Jorgensen et al. 1983). Sodium and chloride ions counterbalanced the charge of the system. The particle-mesh Ewald summation method was used for long-range electrostatics and a 10 Å cutoff was set for short-range non-bonded interactions. Initial geometries in all systems were minimized at 5,000 conjugate-gradient steps after which water was equilibrated at 298 K and 1 atm for 100 ps at 2 fs time steps. Production runs were then performed for 250 ns in the NPT ensemble at P = 1 atm and T = 298 K. We used the hydrogen mass partitioning method to ensure the stability of the simulation under large integration steps, thereby increasing the simulation speed. Langevin dynamics for T control and Nosé-Hoover Langevin piston method for P control were used. We carried out the MD simulation on OpenMM 7.7 (Eastman et al. 2017) running in the Nvidia Tesla A100, L40 and H100 GPU nodes of the Center for High Throughput Computation at the University of Wisconsin-Madison. We performed two replicates of MD simulations. The equilibration of simulations across different complexes was evaluated by tracking the RMSD of the complex structure over time for both replicates (**supplementary fig. 8**). The RMSD across all RuBisCO complexes stabilizes at a plateau, indicating that the complexes reached an equilibrated state.

### Gas diffusion MD simulations

We obtained CO_2_ and O_2_ parameters through the standard Amber protocol. We computed the charges of these molecules at the HF/6-31*G level through the RESP protocol (Woods and Chappelle 2000). The embedding of proteins in solutions with gas molecules in dissolution consisted of replacing water molecules from a previously solvated system to avoid clashes among the gas molecules. We calculated the number of molecules corresponding to a given concentration using the Amber-recommended method for determining ion concentrations in a solvent. All other simulation aspects were consistent with those described in the previous section. These simulations were run for 75 ns, with the final 50 ns used for analysis following a 25 ns equilibration period.

### Analysis of MD simulations

We employed MDAnalysis (Michaud-Agrawal et al. 2011) and ProDy (Bakan et al. 2011) to analyze and process the outcome of the MD simulations. All simulations were conducted under identical temperature, pH, and solvent conditions. The initial 50 ns of the 250 ns simulations were designated for equilibration, and only the final 200 ns were used for analysis. Data from both MD simulation replicates were included, resulting in a combined simulation time exceeding 7 µs. The average system size was approximately 300,000 atoms, including solvent molecules.

### RMSD and RMSF calculations

We aligned all the simulation trajectories to their average structure and calculated the RMSD across all Cα backbone atoms for each RbcL and RbcS subunit using MDAnalysis. The boxplot in Figure 4B presents the average RMSD value for each large and small subunit across the two simulation replicates. RMSF was calculated by aligning the MD trajectories to their reference structure first. The Cα atomic positions of each residue were mapped onto the respective sequence, and fluctuations were calculated as the average positional deviation from the mean structure over the simulation period across the eight RbcL subunits using MDAnalysis. Simulation data from replicates were included in the calculations of the RMSD and RMSF.

### Principal component analysis

The principal component analysis helps reduce the complexity of MD simulation data by generating orthogonal eigenvectors (principal components, PCs) that represent the primary axes of motion. We conducted PC analyses using the scikit-learn implementation (Pedregosa et al. 2011). Specifically, we first extracted all conserved residues across select RbcL sequences shown in **Fig. 4A**. All the individual trajectories for each RbcLs were then aligned onto the reference extant RbcL structure (1BWV). The covariance matrix of positional fluctuations was constructed exclusively for the conserved RbcL Cα atoms. Eigenvalues and eigenvectors were derived from this covariance matrix to identify the PCs shared across all RbcL trajectories, including each simulation replicate. PCA, including the Anc-I without RbcS (**supplementary fig. 10A**), was conducted by extracting trajectories for conserved residues and projecting them onto the PC-space defined by Form-I and Form-I’ RuBisCOs in **Fig. 5**. Additionally, a separate PCA was performed following the same procedure, incorporating RbcL simulation trajectories from Form-II alongside Form-I and Form-I’, as shown in **supplementary fig. 10B**.

## Data and resource availability

The data underlying this article along with the code for analysis can be found at https://github.com/kacarlab/RuBisCO_evolution. Molecular dynamics simulation files for the extant and ancestral RuBisCOs are available at Zenodo (https://zenodo.org/records/14187581).

## Acknowledgements

We would like to thank Alessandro Senes, Phil Romero, Tina Wang and Srivatsan Raman for their feedback on the study, Aya Klos, Evrim Fer, Josh MacCready, Zach Adam and Amanda Garcia for critical reading and useful comments and suggestions on the manuscript. The Center for High Throughput Computing (CHTC) at the University of Wisconsin-Madison provided computing resources for this work. This work was supported by the Human Frontier Science Program (HFSP) [*RGY0072/2021*] with additional support from the NASA Exobiology [*NNH23ZDA001N*] and ICAR Programs [*80NSSC17K0296*]. B.C.Z acknowledges the Margarita Salas Postdoctoral Fellowship, founded by the Unión Europea - Next Generation EU (B.C.Z.; UP2021-035).

## Author Contribution

Conceptualization: KA, BCZ and BK; Methodology: KA and BCZ; Data curation: KA and BCZ; Data analysis: KA, BCZ and BK; Figures: KA and BCZ; First draft: KA; Resources and supervision: BK. All authors edited and approved the final draft.

## Conflict of Interest

The authors declare no competing interests.

## Notes

https://github.com/kacarlab/RuBisCO_evolution

### Competing Interest Statement

The authors have declared no competing interest.

### Summary of Updates

Additional analyses, updated text, a new Fig in the main text, SI files updated.

