## Supplemental Information for "Evolutionary Dynamics of RuBisCO: Emergence of the Small Subunit and its Impact Through Time"

Short Title: Small subunit emergence and RuBisCO evolution

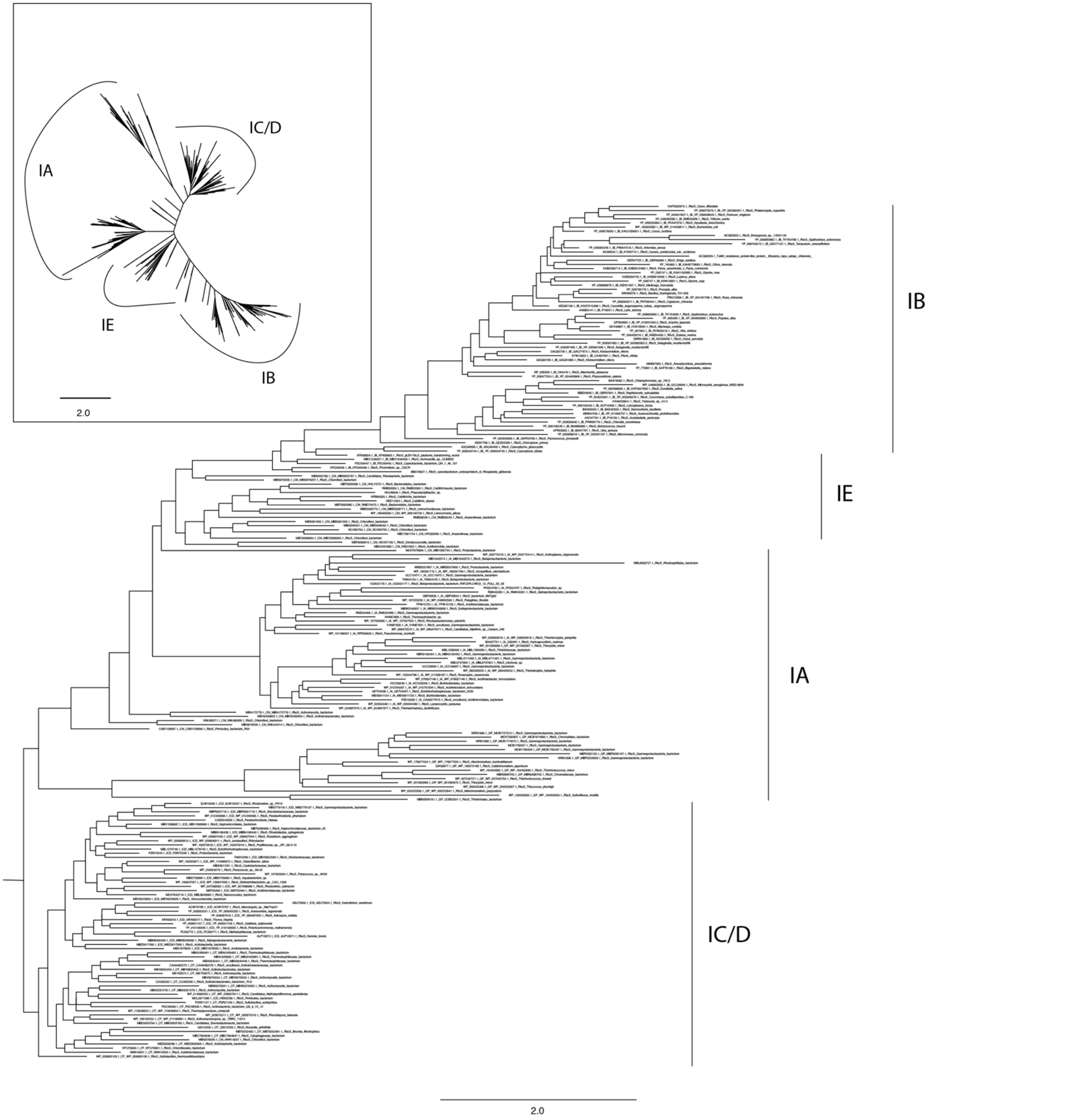


Supplementary Fig. 1: Form-I RuBisCO RbcS Maximum-likelihood rooted and the unrooted phylogeny.

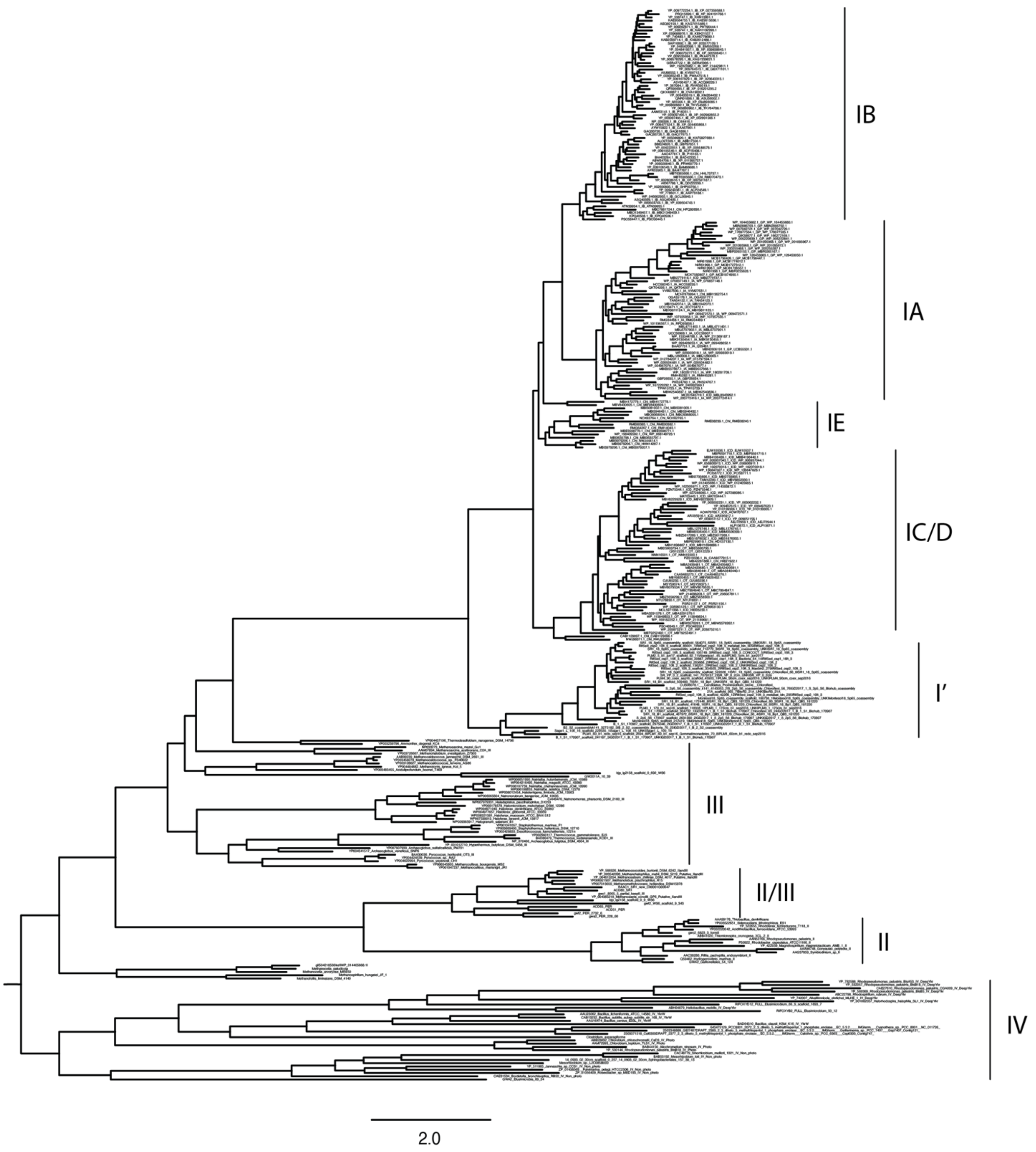


Supplementary Fig. 2: RuBisCO RbcL Maximum-likelihood phylogeny highlighting all RuBisCO Forms.

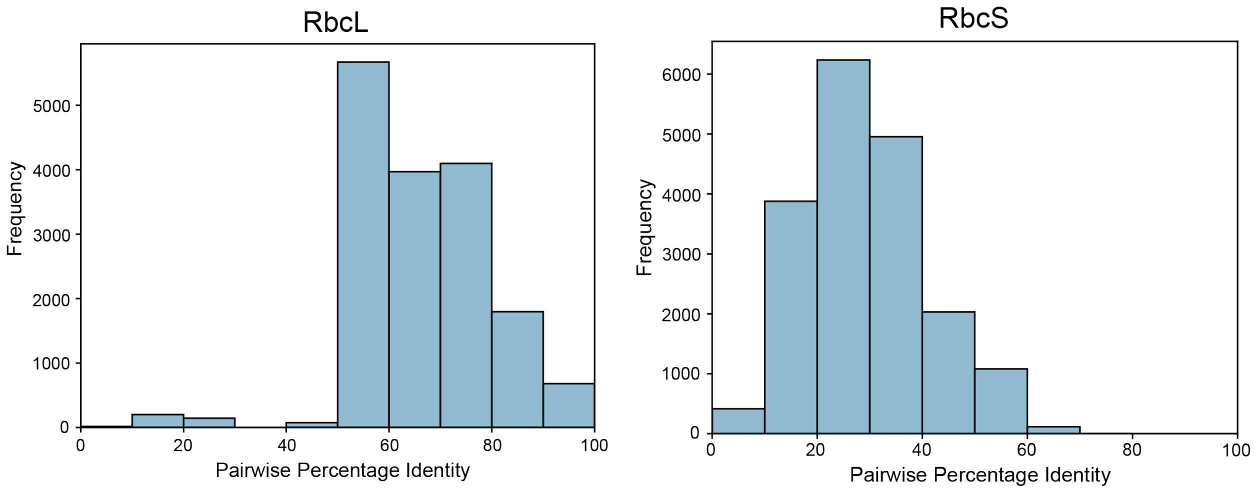


Supplementary Fig. 3: Histogram representing the distribution of the pairwise percentage identity between all Form-I RbcL and RbcS sequences used for building the phylogenetic tree.

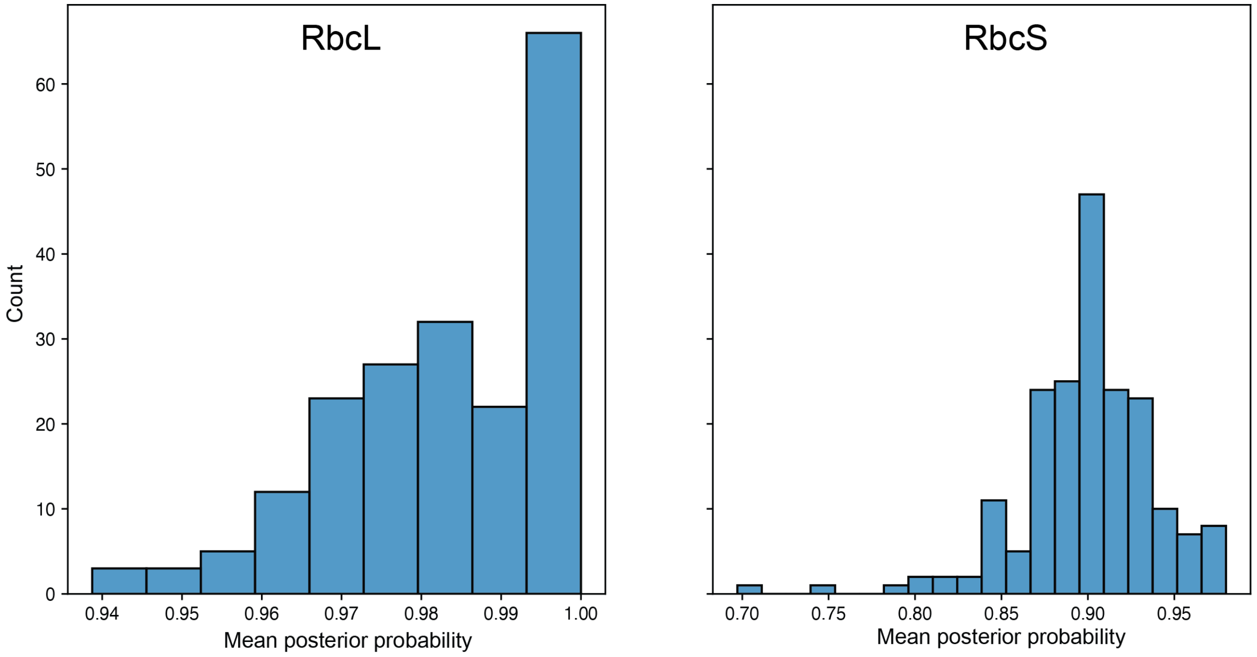


Supplementary Fig. 4: Histogram representing the distribution for mean posterior probability across all positions in the maximum-likelihood Form-I ancestral sequences.

Supplementary Table 1: Variation in sequence features and kinetic properties across Form-I RuBisCOs.

| | **RbcS sequence/structure features** | | | **Biochemical Properties** | | |
|---|---|---|---|---|---|---|
| | **N-terminal Domain** | **βA-βB Loop** | **βE-βF hairpin** | **Specificity, $(K_{cat,C}/K_{M,C})/(K_{cat,O}/K_{M,O})$** | **Carboxylation Rate, $K_{cat,C}$ ($s^{-1}$)** | **Organism** |
| **Form-IB** | - | - | - | 43 | 14.4 | *Synechococcus elongatus* (Cyanobacteria) (Occhialini et al. 2016) |
| | + | + | | 82 | 2.9 | *Spinacia oleracea* (Plantae)(Badger et al. 1998) |
| **Form-IA** | - | - | - | 25.9 | 3.7 | *Rhodobacter capsulatus* (Purple Non Sulfur) (Horken and Tabita 1999) |
| **Form-IE** | - | - | - | 12.2 | 1.1 | *Anaerolineae bacterium* (Chloroflexi) (Schulz et al. 2022) |
| **Form-IC/D** | - | - | + | 114 | 5.7 | *Phaeodactylum tricornutum* (Diatom) (Badger et al. 1998) |
| | | | | 238 | 1.6 | *Galdieria partita* (Red Algae) (Uemura et al. 1997) |

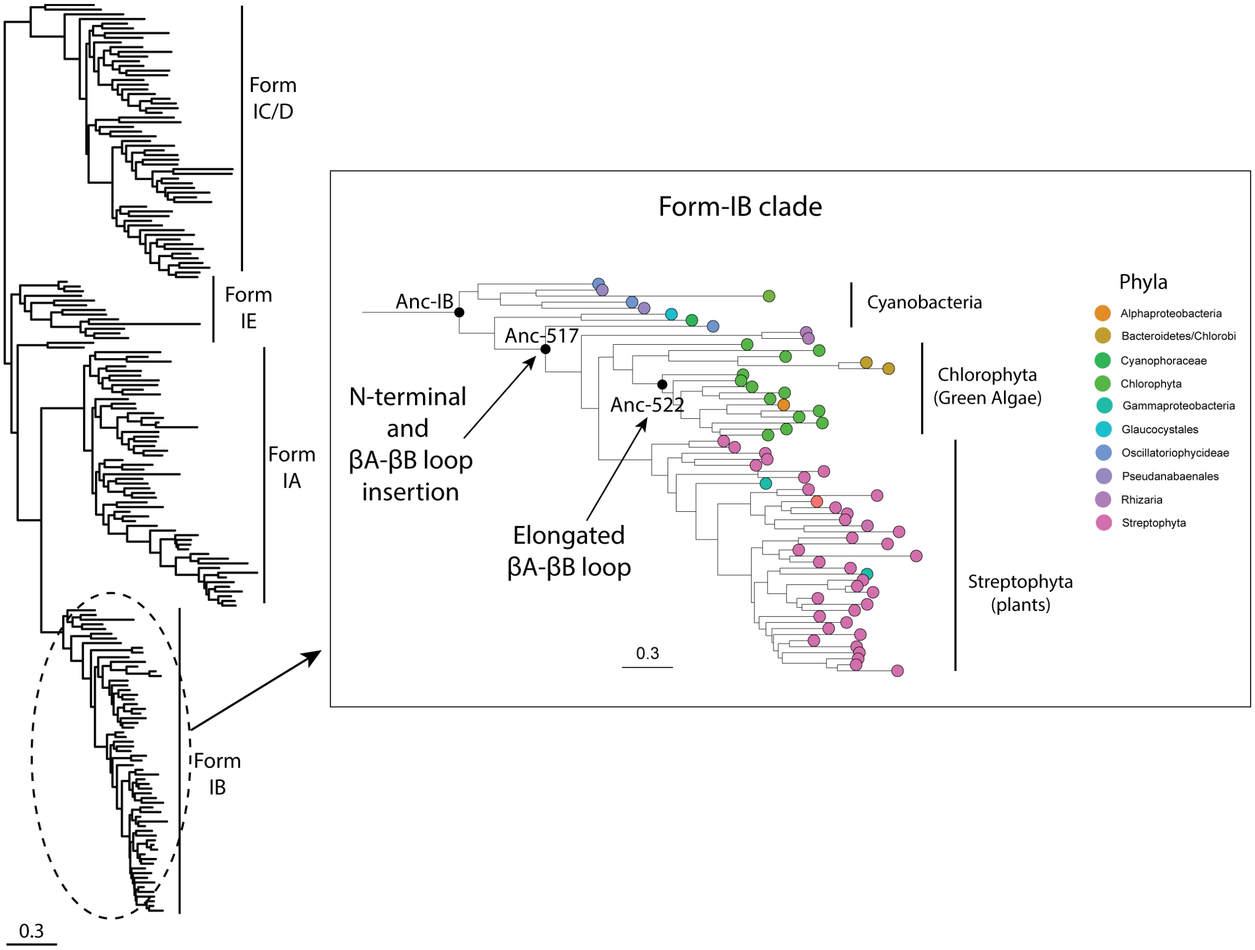


Supplementary Fig. 5: Details for the emergence of different RbcS features within Form-IB RuBisCO structures. Inset displays the detailed Form-IB clade phylogeny highlighting the insertion of N-terminal and βA-βB loop in RbcS.

Supplementary Table 2: Emergence of different RbcS structural features along RuBisCO's evolutionary trajectory.

| **Feature** | **Ancestral node of first emergence** | **Information about the ancestor** | **Functional Relevance** |
|---|---|---|---|
| N-terminal extension | Anc-517 | First ancestor after divergence from cyanobacteria in Form-IB clade phylogeny. | Signal peptide necessary for the entry of RbcS into the chloroplast (Schmidt and Mishkind 1986). |
| βA-βB loop insertion | Anc-517 | | Functional role not known; regulates size of the central solvent channel in the RuBisCO complex (Esquivel et al. 2013). |
| Extension of βA-βB loop insertion | Anc-522 | Common ancestor for the majority of green algae species in the phylogeny of Form-IB clade. | Same as above; Extension of the loop is predominantly found in RbcS from green algae (Spreitzer 2003). |
| βE and βF hairpin loop | Anc-IC/D | Common ancestor of all Form-IC/D RuBisCOs. | Mediates the assembly of the RuBisCO oligomeric complex in red-like RuBisCOs (Joshi et al. 2015). |

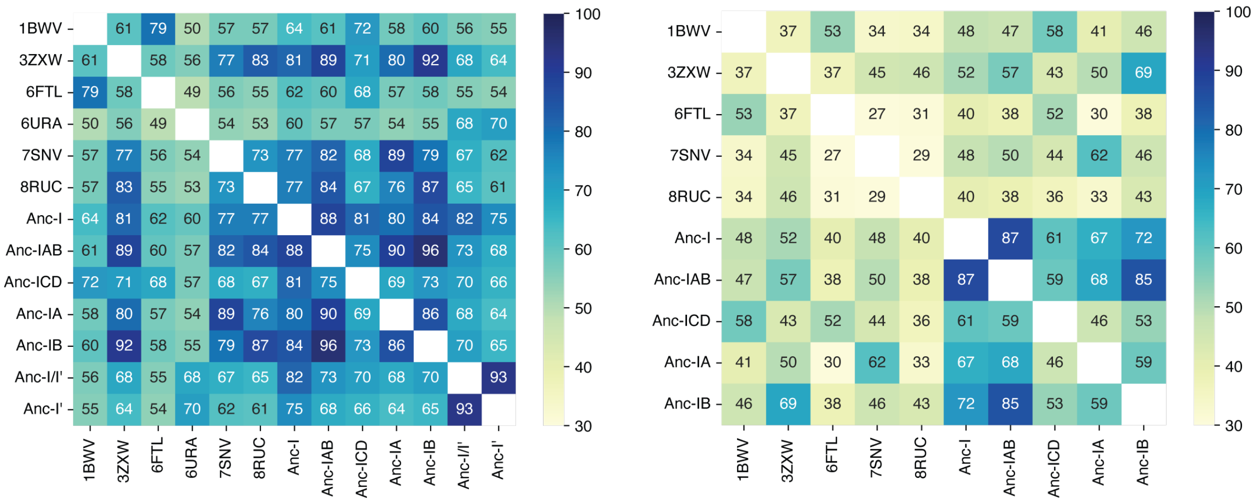


Supplementary Fig. 6: Sequence identity matrix for the ancestral and extant RuBisCO large (left) and small (right) subunit used for the MD-simulations.

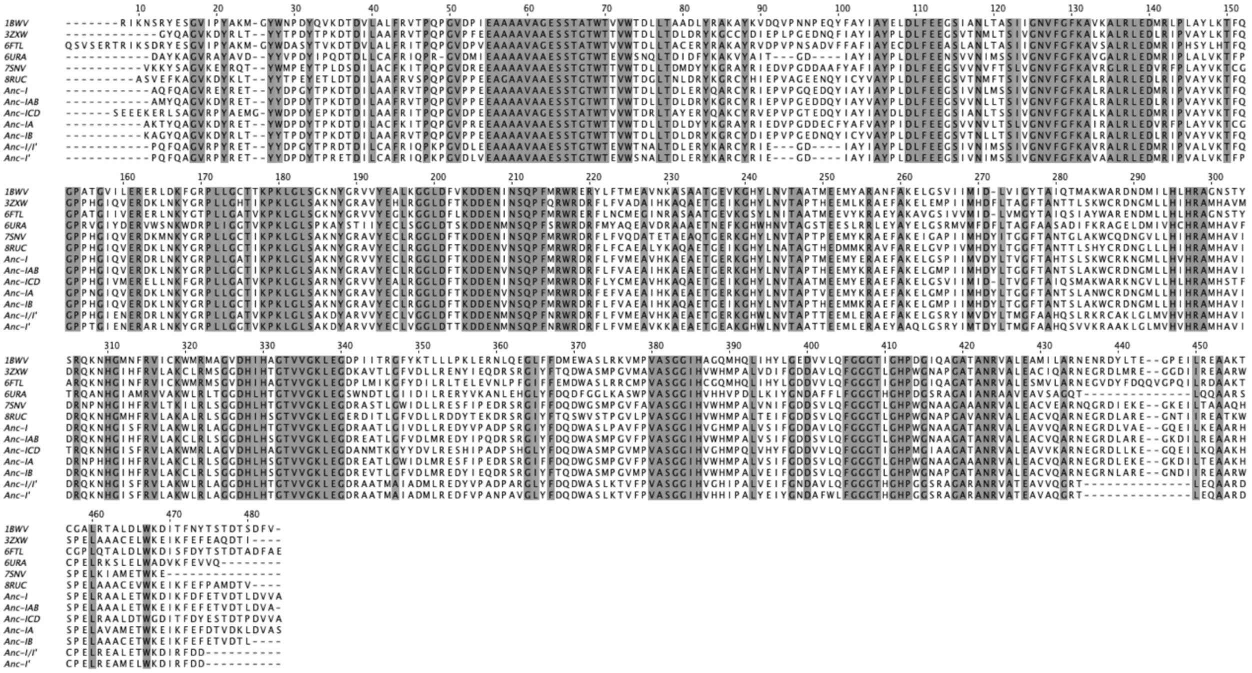

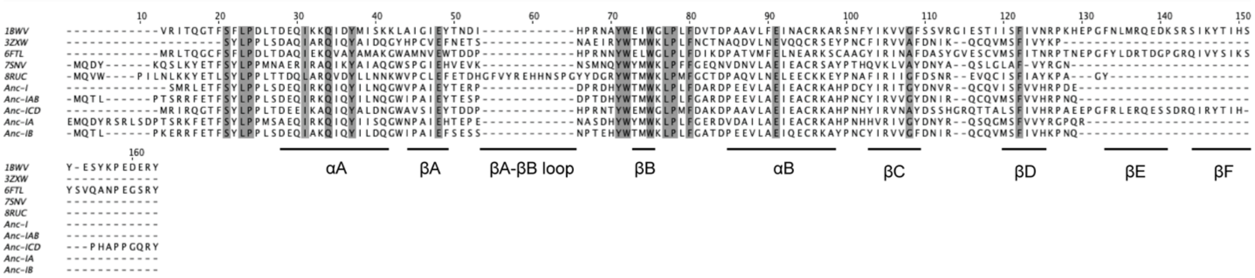


Supplementary Fig. 7: Multiple sequence alignment for the ancestral and extant RuBisCO large (top) and small (bottom) subunit used for the MD simulations.

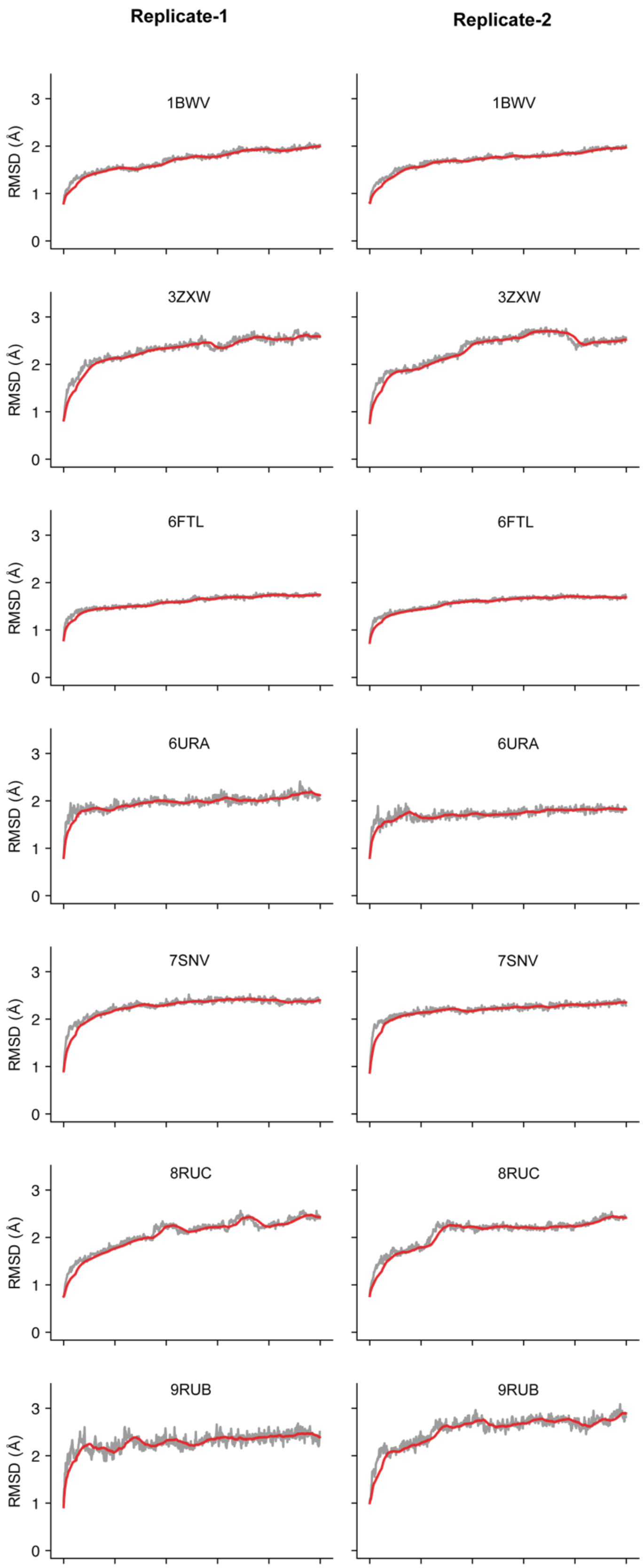
Replicate-1
Replicate-2
1BWV
3ZXW
6FTL
6URA
7SNV
8RUC
9RUB
RMSD (Å)
0
1
2
3

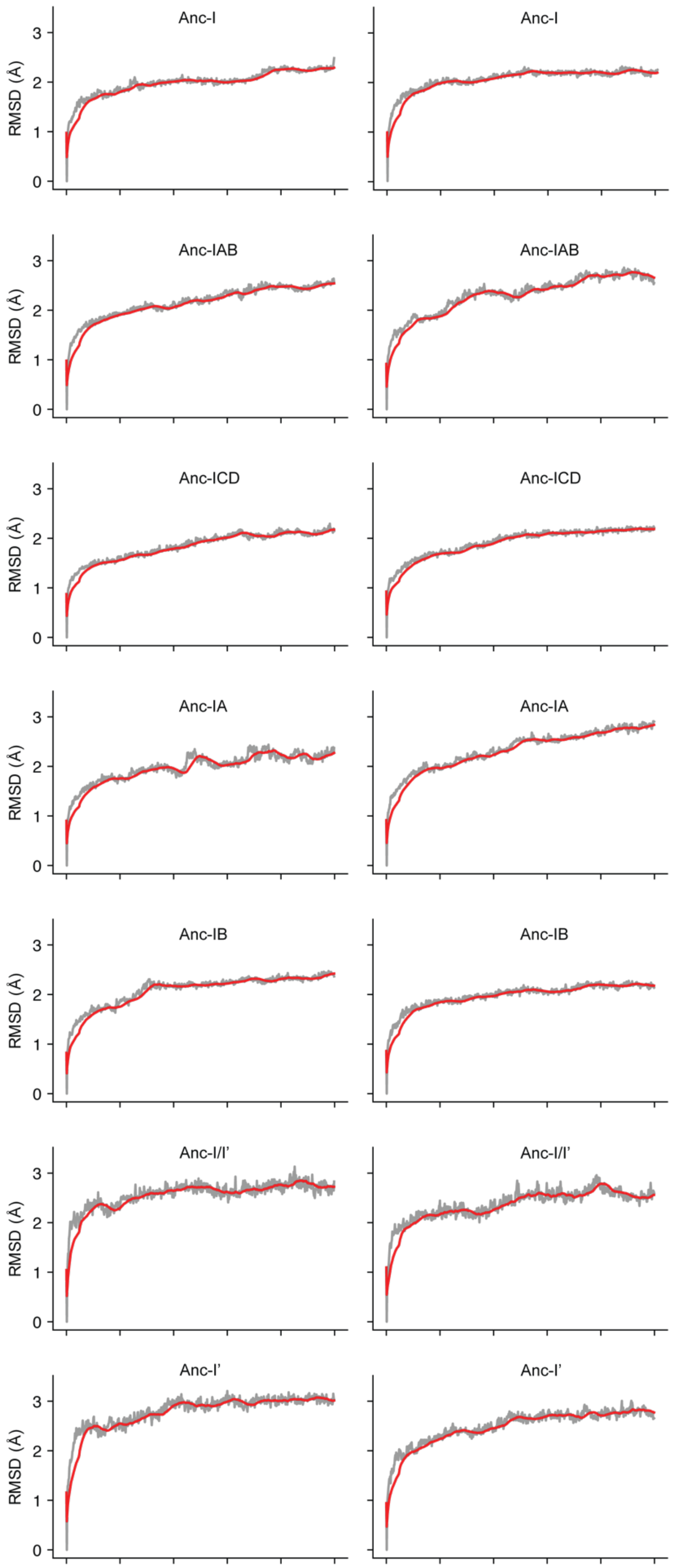
Anc-I
Anc-I
Anc-IAB
Anc-IAB
Anc-ICD
Anc-ICD
Anc-IA
Anc-IA
Anc-IB
Anc-IB
Anc-I/I'
Anc-I/I'
Anc-I'
Anc-I'
RMSD (Å)
0
1
2
3

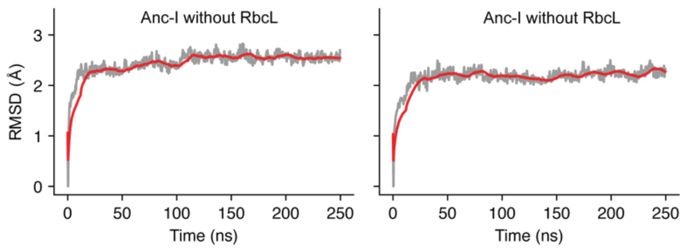


Supplementary Fig. 8: The root mean square deviation (RMSD) over time for all Cα-atoms in the complex across ancestral and extant RuBisCOs during the MD for 2 simulation replicates. The left and right column represents the values for the first replicate and second replicate respectively. The shaded gray line represents raw RMSD data, while the red solid line shows values averaged over a 30 ns window.

Supplementary Table 3: RuBisCO pairs with significant ($p < 0.01$) difference in mean RMSD for RbcL during the MD-simulations.

| **RbcL pairs** | **Difference in mean RMSD (in Å)** | **p-value (Two-sided)** |
|---|---|---|
| 1BWV – Anc-I’ | 0.2929 | 0.0003 |
| 3ZXW – 6FTL | -0.2473 | 0.0063 |
| 6FTL – Anc-I’ | 0.3766 | 0.0000 |
| 6FTL – Anc-I/I’ | 0.2873 | 0.0005 |
| 6FTL – Anc-IA | 0.2516 | 0.0049 |
| 6URA – Anc-I’ | 0.3056 | 0.0001 |
| 7SNV – Anc-I’ | 0.3523 | 0.0000 |
| 7SNV – Anc-I/I’ | 0.2630 | 0.0025 |
| Anc-I – Anc-I’ | 0.2420 | 0.0086 |
| Anc-IB – Anc-I’ | 0.2464 | 0.0067 |

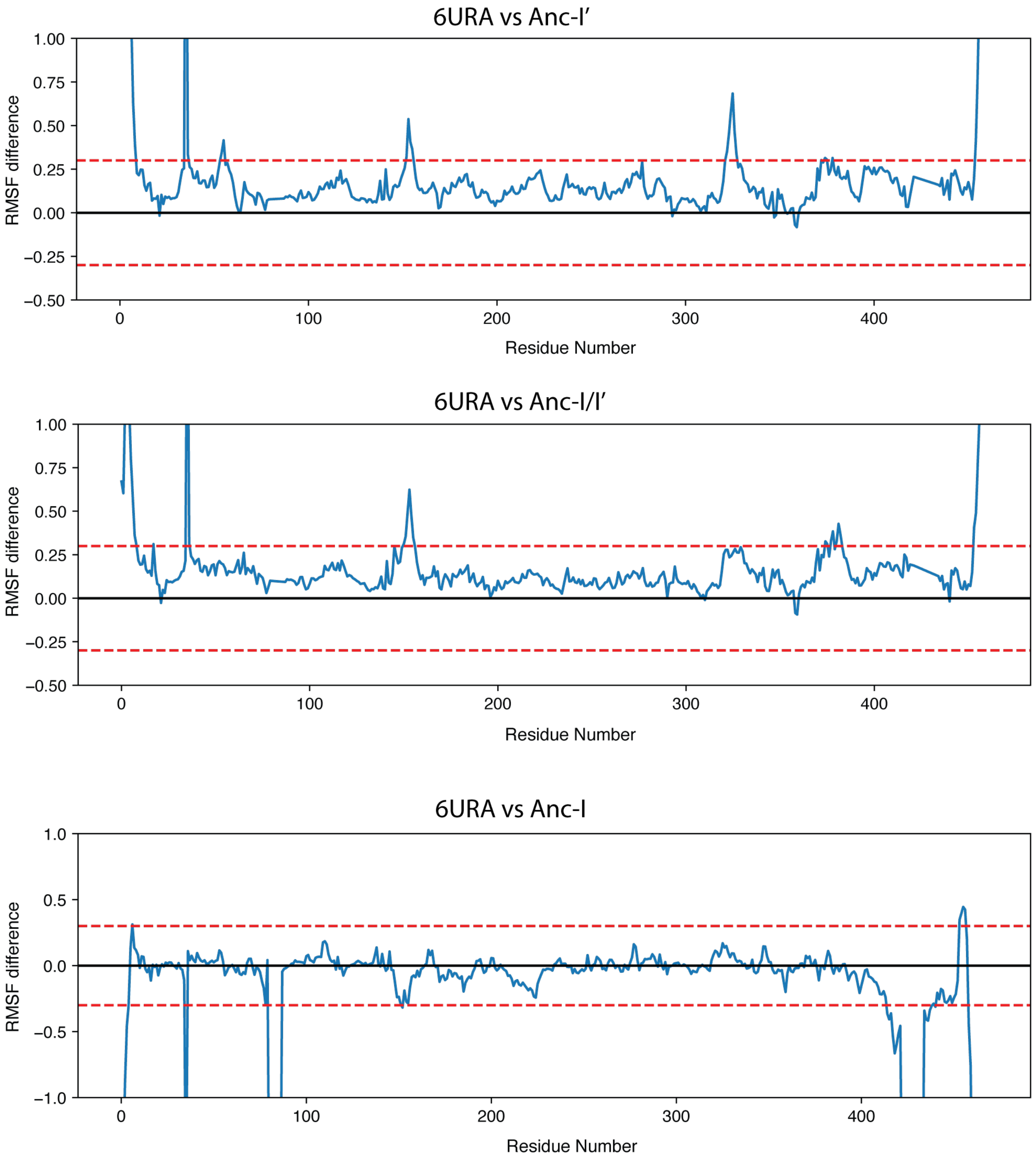


Supplementary Fig. 9: Differences in residue-wise RMSF for the 6URA RbcL with Anc-I’, Anc-I/I’ and Anc-I. The first two plots show that Anc-I’ and Anc-I/I’ display higher structural fluctuations compared to 6URA across the length of RbcL over the course of simulation.

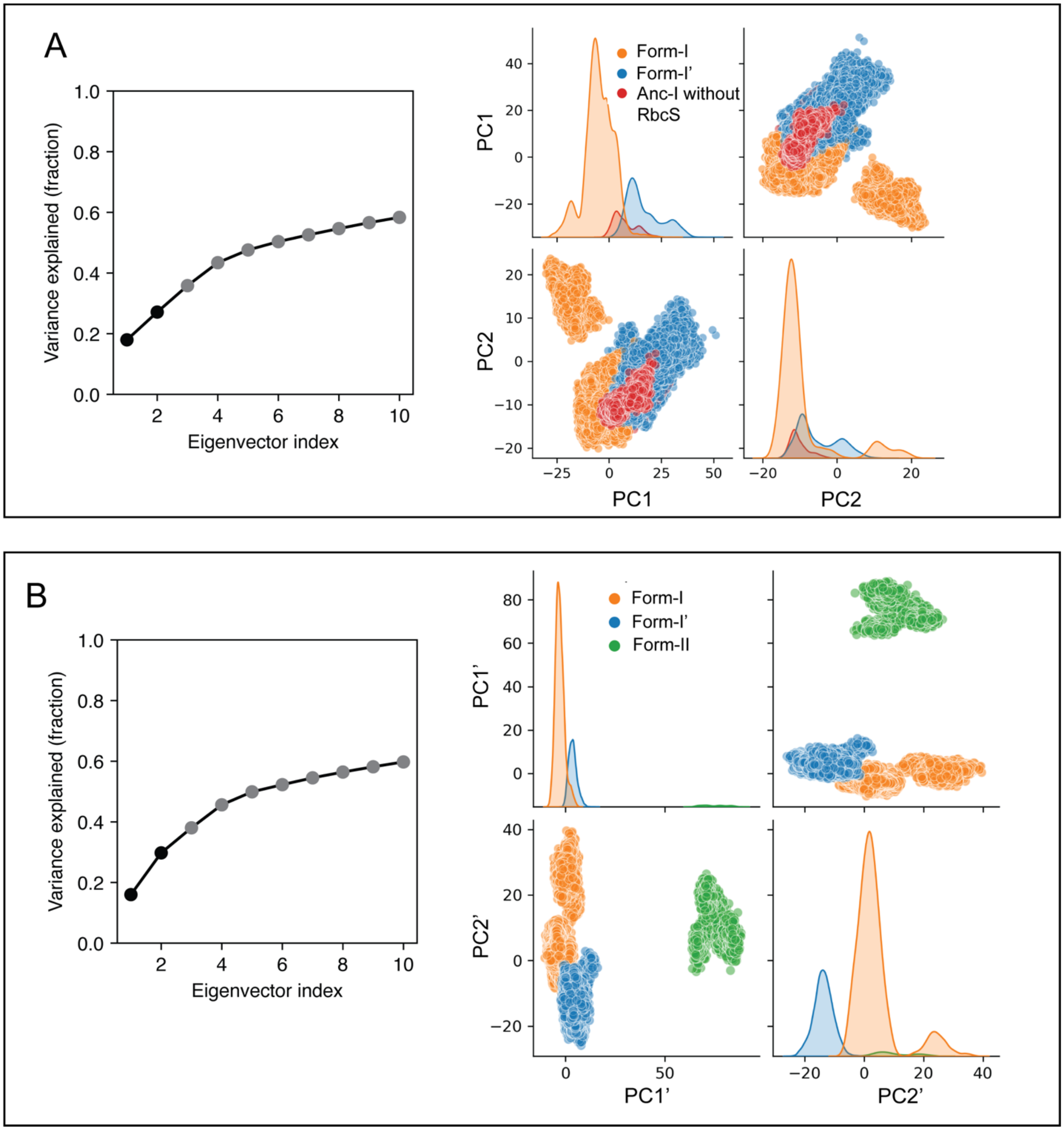


Supplementary Fig. 10: Principal Component Analysis of RbcL residues including the Form-I without RbcS and Form-II RuBisCO variants. (A) Variance explained by each eigenvector in the PCA and pairwise representation of PC1 and PC2 after mapping the trajectory for Anc-I RuBisCO simulations without RbcS onto the PCs for Form-I and Form-I' RuBisCO. (B) Variance explained by each eigenvector in the PCA and pairwise representation of PC1' and PC2' for Form-I, Form-I' and Form-II RuBisCOs.

Supplementary Table 4: Values for the Gas Contact Analysis for different MD-simulations.

| **System** | **Subunit** | **$CO_2$** | **$O_2$** | **t-statistic** | **p-value (two-sided)** |
|---|---|---|---|---|---|
| 1BWV | RbcL | 151.2 ± 29.64 | 80.34 ± 11.17 | 5.918860 | 3.743741e-05 |
| 1BWV | RbcS | 224.34 ± 26.71 | 107.46 ± 23.29 | 8.726068 | 4.911744e-07 |
| 3ZXW | RbcL | 92.1 ± 10.18 | 90.17 ± 8.44 | 0.385388 | 7.057424e-01 |
| 3ZXW | RbcS | 161.25 ± 35.65 | 121.45 ± 31.33 | 2.218684 | 4.354644e-02 |
| 6FTL | RbcL | 113.44 ± 43.27 | 88.36 ± 29.39 | 1.268552 | 2.252889e-01 |
| 6FTL | RbcS | 94.81 ± 22.33 | 95.86 ± 33.27 | -0.069357 | 9.456860e-01 |
| 7SNV | RbcL | 81.59 ± 7.56 | 73.59 ± 6.85 | 2.073243 | 5.708508e-02 |
| 7SNV | RbcS | 158.2 ± 22.22 | 111.75 ± 10.95 | 4.960129 | 2.094638e-04 |
| 8RUC | RbcL | 107.19 ± 31.06 | 87.55 ± 21.78 | 1.370376 | 1.921438e-01 |
| 8RUC | RbcS | 111.57 ± 35.01 | 101.81 ± 20.99 | 0.632452 | 5.372848e-01 |
| Anc-I | RbcL | 93.75 ± 16.1 | 92.86 ± 13.94 | 0.110304 | 9.137338e-01 |
| Anc-I | RbcS | 136.6 ± 18.35 | 109.67 ± 21.78 | 2.501628 | 2.538669e-02 |
| Anc-IA | RbcL | 94.82 ± 19.25 | 75.59 ± 16.5 | 2.007307 | 6.442361e-02 |
| Anc-IA | RbcS | 151.66 ± 32.79 | 101.9 ± 21.59 | 3.352923 | 4.736344e-03 |
| Anc-IAB | RbcL | 96.08 ± 14.98 | 94.06 ± 16.32 | 0.241688 | 8.125261e-01 |

| **System** | **Subunit** | **$CO_2$** | **$O_2$** | **t-statistic** | **p-value (two-sided)** |
|---|---|---|---|---|---|
| Anc-IAB | RbcS | 142.66 ± 21.17 | 98.23 ± 17.23 | 4.305906 | 7.252118e-04 |
| Anc-IB | RbcL | 87.27 ± 9.11 | 86.22 ± 6.51 | 0.247780 | 8.079016e-01 |
| Anc-IB | RbcS | 115.03 ± 13.94 | 108.72 ± 18.54 | 0.719665 | 4.835733e-01 |
| Anc-ICD | RbcL | 73.71 ± 13.33 | 76.88 ± 9.26 | -0.516770 | 6.133833e-01 |
| Anc-ICD | RbcS | 135.56 ± 30.9 | 130.16 ± 18.08 | 0.398578 | 6.962165e-01 |

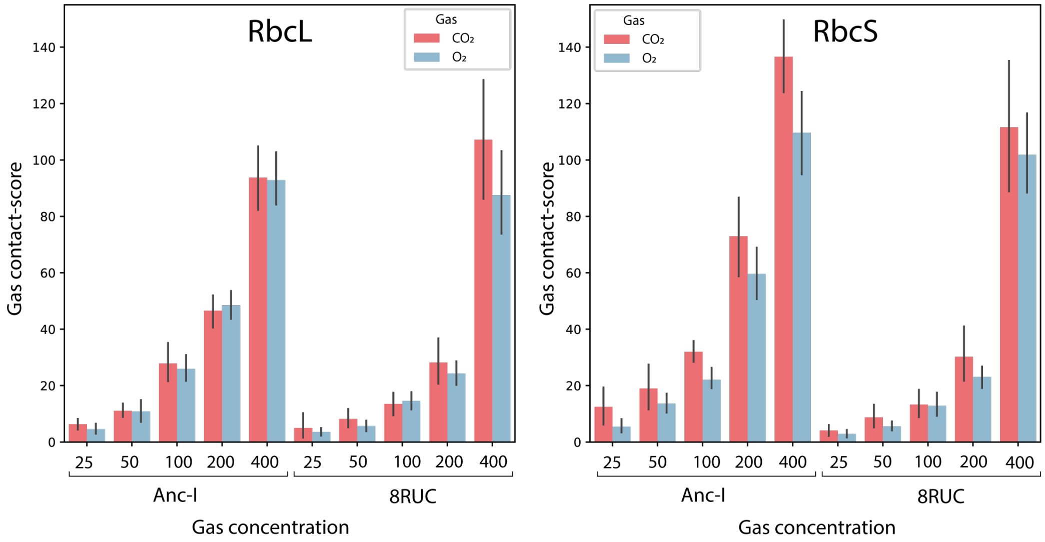


Supplementary Fig. 11: $CO_2$ and $O_2$ gas contact-score for RbcL and RbcS for the Anc-I and 8RUC RuBisCO systems with different gas concentrations.

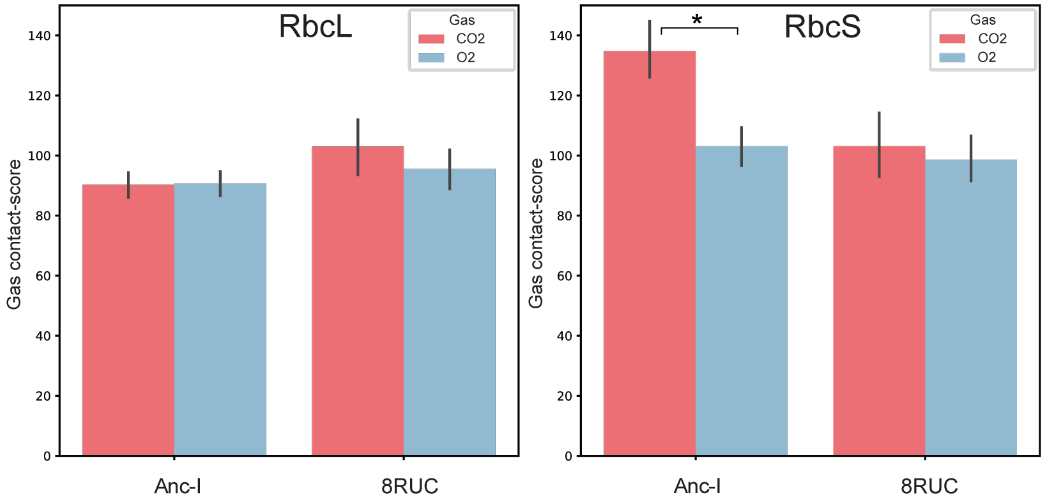


Supplementary Fig. 12: $CO_2$ and $O_2$ gas contact-scores for RbcL and RbcS for the Anc-I and 8RUC RuBisCO systems over 5 replicates.
